# Cerebellar associative learning underlies skilled reach adaptation

**DOI:** 10.1101/2021.12.17.473247

**Authors:** Dylan J. Calame, Matthew I. Becker, Abigail L. Person

## Abstract

Cerebellar output has been shown to enhance movement precision by scaling the decelerative phase of reaching movements in mice. We hypothesized that during reach, initial kinematics cue late-phase adjustments through cerebellar associative learning. We identify a population-level response in mouse PCs that scales inversely with reach velocity, suggesting a candidate mechanism for anticipatory control to target limb endpoint. We next interrogate how such a response is generated by combining high-density neural recordings with closed-loop optogenetic stimulation of cerebellar mossy fiber afferents originating in the pontine nuclei during reach, using perturbation schedules reminiscent of classic adaptation paradigms. We found that reach kinematics and PC electrophysiology adapt to position-locked mossy fiber perturbations and exhibit aftereffects when stimulation is removed. Surprisingly, we observed partial adaptation to position-randomized stimulation schedules but no opposing aftereffect. A model that recapitulated these findings provided novel insight into how the cerebellum deciphers cause-and-effect relationships to adapt.

## Introduction

In humans and animals with altered cerebellar function, movement is disorganized, often showing hallmark symptoms of endpoint dysmetria^1,2^ and impaired abilities to adapt movements in the face of novel conditions^3–7^. These observations have led to the idea that the cerebellum makes movements fast, smooth, and accurate by learning anticipatory control signals that mediate feedforward control ^8–10,11,12^.

Understanding the neurobiological basis of learned anticipatory signals is a major effort of cerebellar physiology. Two dominant learning paradigms, classical conditioning and motor adaptation, each provide important insight into the mechanisms of anticipatory motor control. Classical conditioning paradigms^13–15^ such as delay eyeblink conditioning, illustrate how neutral conditioned stimuli paired with reflex-eliciting unconditioned stimuli become predictive cues eliciting conditioned responses (e.g. a tone repeatedly paired with a corneal airpuff eventually elicits a predictive eyeblink). Mechanistically, neutral cues can be fully replaced by cerebellar mossy fiber stimulation^16,17^ and unconditioned stimuli can be fully replaced by climbing fiber stimulation^17^. Climbing fibers elicit complex spikes (Cspks) in Purkinje cell (PC) dendrites^18,19^, which over many trials reduce parallel fiber efficacy onto PCs, leading to firing rate pauses at the predicted time of the unconditioned stimulus. Through subsequent disinhibition of the cerebellar nuclei, these pauses then drive anticipatory conditioned responses.

Cerebellar cortex also mediates many forms of motor adaptation^20–22^. Perturbations that result in movement error, such as prisms that distort gaze or target jumps that cause endpoint errors, modulate Cspks to induce bidirectional plasticity in PC simple spike activity that correlates with adaptive changes in behavior^23–25^. It has been hypothesized that such instances of motor adaptation can be understood through a lens of associative learning. In this view, sensorimotor information, conveyed to the cerebellum via mossy fibers, is reweighted onto Purkinje neurons when associated with deviations from basal climbing fiber rates. Learning mechanisms analogous to those described above for associative conditioning then reweight the original sensorimotor inputs to Purkinje neurons leading to novel cerebellar output^25^.

Despite the widely held view that motor adaptation shares a mechanistic substrate with classical conditioning, key differences between motor control and classical conditioning make these inferences tenuous. For example, perturbations that drive motor adaptation^26–28^ engage sensorimotor feedback loops at multiple levels of the nervous system, complicating the view that cerebellar input-output remapping fully explains adaptation since inconstant cerebellar inputs would deprive associative mechanism of a stable cue. This problem is even more exaggerated when considering skilled movements such as reach, where both cerebral cortex and cerebellum are proposed as sites of learning^26,29,30^, with long-range loops likely functioning synergistically^31^. Thus, learning-related changes in cerebellar Purkinje firing could be inherited from cerebral cortex, generated by cerebellar plasticity, or both^32^.

We hypothesize that cerebellar output during movements is mechanistically akin to conditioned responses, such that in learned conditions, cerebellar outputs reduce motor error and are cued by mossy fiber activity that encodes within-movement kinematic features^10,25^. Recent studies in mice performing skilled reach movements suggest experimental entry points to testing this idea. Specifically, a population of cerebellar output neuron firing rates scale with and cause reach deceleration, and covary with early kinematic features like peak reach velocity^2,33^. Moreover, acute disruption of cerebral cortical input to the cerebellum impairs skilled reach kinematics^2,34,35^, suggesting that cerebellar inputs could be used both as a perturbation and a cue to predictively adjust skilled movement kinematics.

Here, we performed experiments designed to link the cerebellum’s role in associative learning with its generation of anticipatory control of reaching movements. We superimposed a variant of associative learning onto a skilled reach task through repeated optogenetic manipulation of mossy fiber activity in closed loop with reach, triggered at a consistent kinematic landmark. In addition to monitoring the kinematic effect of stimulation over trials, we also measured changes in PC response to stimulation. To test reliance of temporal specificity on learning, we randomized the position of stimulation. A cerebellar model of timed adaptation within a movement recapitulated our key experimental findings and gives mechanistic insight into the circuit properties underlying cerebellar reach adaptation. Together, these experiments unify the frameworks of cerebellar associative learning and motor adaptation in skilled movements, an important step in understanding mechanisms of motor learning.

## Results

### A suppression-based Purkinje cell population code tuned to reach velocity

Neurons in the anterior interposed nucleus fire proportional to reach velocity and causally scale limb deceleration, such that the limb lands on target despite initial kinematic variability^2,33^, consistent with the cerebellum implementing anticipatory control. To determine whether upstream PCs may drive these decelerative bursts in the cerebellar nuclei, we combined kinematic and electrophysiological recordings in mice engaged in a skilled head-fixed reach task. After mice were proficient at the task, we recorded reach kinematics with high-speed cameras via an IR-reflective marker affixed to the mouse’s hand (Fig. 1a, Fig. S1). Acute recordings in cerebellar cortex were made simultaneously, using either single electrodes or Neuropixel probes (Fig. 1a, Video S1). Recordings were targeted to a cerebellar cortical site situated between Lob 4/5 and Simplex known to influence forelimb movements in mice^13^. Putative PCs were identified as neurons with Cspks or with a firing rate > 40 spikes/s, CV2 > 0.20, MAD < 0.008 (see Methods, Fig. S2). We found that activity in many PCs was highly modulated around the time of the reach across cells and sessions (Fig 1).

**Figure 1.**
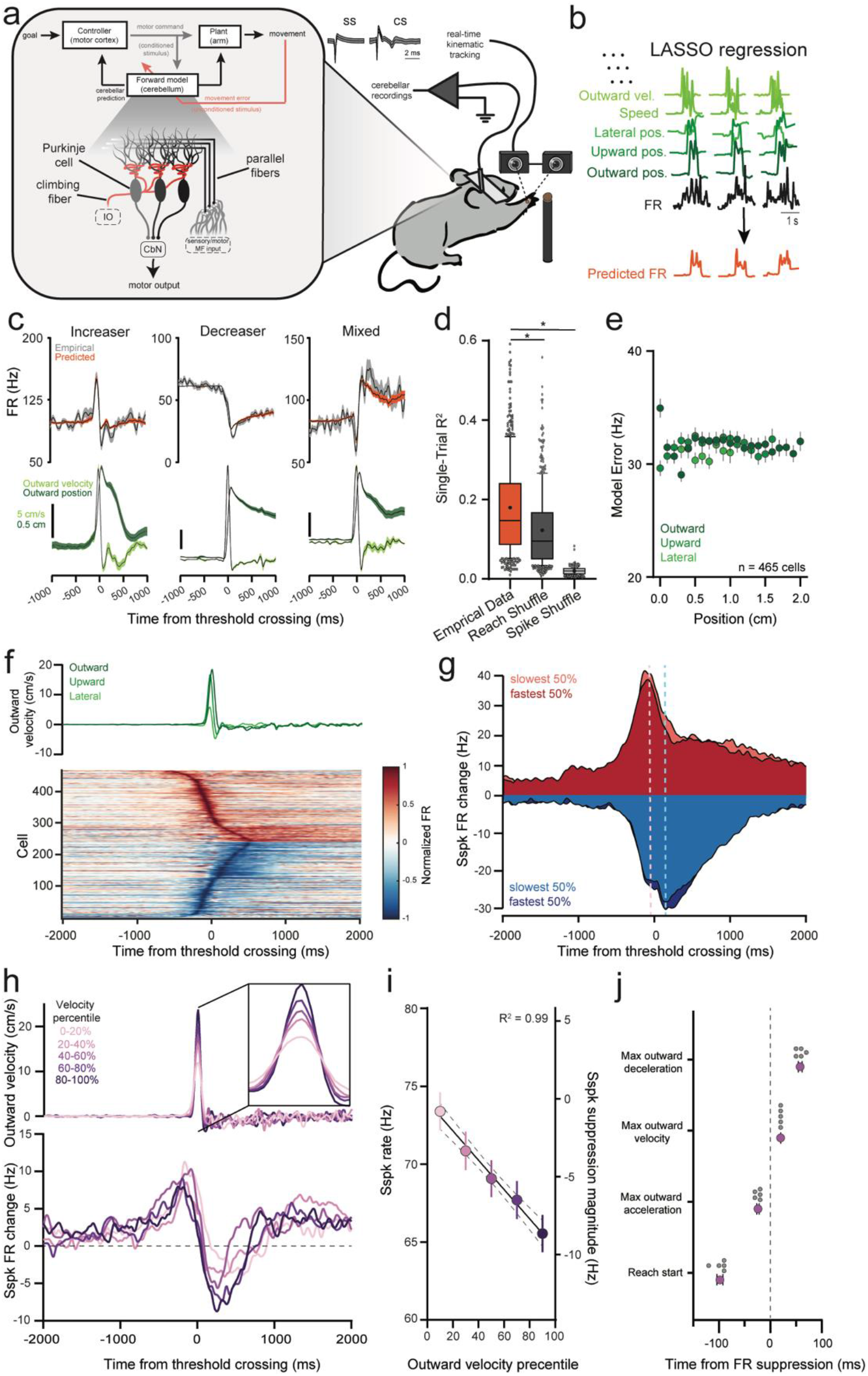
Net population activity in Purkinje cells predicts reach velocity. **a.** Schematic diagram of conceptual framework and experimental paradigm. Predictions computed by the cerebellum are hypothesized to be learned through reweighting of cerebellar inputs, including copies of motor commands, instructed by changes in climbing fiber activity. PCs of the deep central sulcus were recorded with either single electrodes or Neuropixel probes while the reaching hand was tracked in real-time with high-speed cameras. **b.** Kinematic regressors in multilinear LASSO regression were used to model firing rates on individual reaches across sessions. **c.** Examples of 3 PCs fit with LASSO regression. Top: trial-averaged empirical and LASSO predicted firing rates. Bottom: outward position and velocity aligned to firing rate at optimal lag (mean±SEM). **d.** Modest single-trial R^2^ for single cells in empirical, reach shuffled, and spike shuffled LASSO regressions (Box plot shows median line +/− 25%tiles, center dot is mean, whiskers are 10-90%). **e.** Absolute model error (empirical vs predicted, across outward, upward and lateral positions) as a function of reach position. Stable error suggests continuous encoding of reach kinematics across reach epoch. Positions binned at 0.1 cm. **f.** During reach (kinematics, green), PCs group roughly into cells that increase firing rate (red) and cells that decrease firing rate (blue), aligned to the time the hand passed ‘threshold’, 1-cm in the outward direction. **g.** Simple spike firing rate modulation during reaches grouped by reach speed. Cells that increase (red) and decrease (blue) firing rate both showed lower firing rates during faster reaches. **h.** Pooling all PCs reveals net firing rate suppression that scales with reach velocity percentile. Top: Binned reach velocities associated with recordings. Bottom: Net PC population firing rate change for each reach velocity bin. **i.** Magnitude of net firing rate suppression in total PC population as a function of outward velocity. Firing rate during the suppression in population activity was strongly negatively related to reach velocity (mean±SEM). **j.** Time of population suppression is intermediate between peak outward acceleration and peak outward velocity, preceding deceleration. Plot relates the median timing of reach start, peak outward acceleration, peak outward velocity, and peak outward deceleration to the time of population simple spike suppression for each reach velocity bin shown in i (mean±SEM).

To test the prediction that decelerative signals in the cerebellar nuclei derive from Purkinje neuron activity patterns during reach, we first sought to understand what individual PCs encode. We used least absolute shrinkage and selection operator (LASSO) regression to model PC simple spike firing rate using limb kinematics on a trial-by-trial basis, with a ten-fold cross validation step to avoid over-fitting (Fig. 1b; see Methods) ^36^. On average, kinematics of the limb alone could explain a modest 18.0 ± 0.01% (mean ± SEM) of the variance in simple spike firing rate on individual trials, although trial-averaged data was a much closer fit (58.0 ± 0.01%, Fig. 1c, d) consistent with other studies of PC simple spike tuning to limb movements in primates^37–41^. Kinematic encoding was not a result of generic movement-related modulation but was specific to the kinematics of individual reaches as demonstrated by a reach shuffled control that reassigned reaches with PC firing recorded during separate reaches, and a spike shuffled control, where simple spike times on each trial were time shuffled and regressed against kinematics. In both cases, regression performance on the empirical data was significantly higher than the shuffled controls indicating that simple spike firing rates encode reach kinematics on a reach-by-reach basis (Fig. 1d; N = 11 animals, n = 465 cells; empirical vs. reach shuffle: p = 1.1 × 10^−71^, W = 103803, r = 0.83; empirical vs spike shuffle: p = 6.9 × 10^−78^, W = 108331, r = 0.87, Wilcoxon signed rank test). The regression model performance was stable across the spatial trajectory of reaches, suggesting kinematic encoding is continuous in individual cells (Fig. 1e). To assess which kinematic variables in the regression model were the most important in modeling simple spike firing rate, we repeated the regression with each variable independently time shuffled and measured the change in variance explained relative to the complete model^42^ (Fig. S3d). Positional terms -- outward, upward, and lateral – each accounted for approximately 10% of the explained variance of the complete model, with each of the remaining 20 variables accounting for < 5%, although there was a wide variety in the relative importance of different kinematic variables across cells. These measurements are roughly consistent with PC-limb kinematic relationships observed in primates^40,43,44^, however the relatively weak encoding on individual trials obscures how the cerebellum might influence control over movements to make them smooth and accurate.

As has been noted previously, PC simple spike rates fluctuate during movements, with modulations that are either predominately positive or negative ^45,46^. Of 465 PCs recorded during reach, 226 displayed increases in activity during the reach epoch and 239 showed a decrease (Fig. 1f). When segregated into groups that predominantly increase or decrease rates during reach, both populations had lower firing rates during faster reaches relative to slower reaches (Fig. 1g; highest peak velocity). The peak rates of the population of positively modulated cells preceded negatively modulated cells (dashed vertical lines, ~ 220 ms difference), raising the question of how these subpopulations collaborate as a group.

Populations of ~40 PCs converge onto single nuclear cells^47^. In the oculomotor vermis – where heterogenous rate modulation profiles of PCs strongly resemble the patterns we saw during reach – grouping PCs into populations across classes revealed much stronger kinematic relationships with saccades^45,48^. Speculating that similar population encoding principles may be seen in reach-related PCs, we next grouped all PCs across all animals and looked at average activity for reaches binned by outward velocity. Firing rate increases were followed by sharp drops in net activity during the reach epoch that scaled with the velocity of outreach (Fig. 1h). Quantifying the minimum simple spike firing rate during the reach window (see Methods) showed a strong negative relationship with outreach velocity, such that the population showed a suppression of activity that scaled with reach velocity (Fig. 1i; N = 11 animals, n = 465 cells, 2100 reaches; 11806 reach-cell pairs; R^2^ = 0.99, slope = −0.094 with 95% CI [−0.111, −0.077], p = 4.0 × 10^−4^, F = 307.5). The timing of rate suppression was intermediate between peak outward acceleration and peak outward velocity, just before the transition to the decelerative phase of reach (Fig. 1j). Notably, each velocity percentile contained equal populations of positively and negatively modulated neurons. These data suggest that PC suppression scales in a way that is the inverse of decelerative nuclear bursts that causally slow the limb.

Summarizing, we found that individual PCs are privately and modestly tuned to specific kinematic features of reach but weakly related to previously observed patterns of firing in the cerebellar nuclei. Yet, at the population level, PC activity shows scaled suppression in activity shortly before deceleration, consistent with a disinhibitory mechanism driving decelerative bursts in nuclear cells. We hypothesize that this firing rate suppression may be mechanistically akin to conditioned responses seen in delay eyeblink conditioning – learned rate changes that produce anticipatory movements in response to predictive cues. Both the precise timing and scaling of the population activity suppression observed here are consistent with learned cerebellar responses linked to motor and sensory contingencies to control movement. As such, this behavior offers a unique opportunity to test theories relating motor adaptation to associative learning in service of skilled movement^24,49–51^.

### Cspks signal movement onset and reach outcome

To probe mechanisms that might shape cerebellar cortex scaling of output as a function of kinematics, we first identified cerebellar recordings with Cspks, the drivers of learning in PCs. Cspks could be sorted stably across the experiment in 59 of 465 putative PCs, 58 of which had Cspks during the peri-reach window (~1s window centered on reach; see Methods). Cspk probability increased shortly before movement onset, consistent with reports of early synchronized Cspk activity occurring at movement initiation^52–54^, then dropped near steady-state levels (Fig. 2a; p = 3.3 × 10^−5^, F = 2.6, RM one-way ANOVA; mean Cpsk probability vs. −500 ms bin: p = 7.1 × 10^−4^, d = −0.62, Dunnett’s multiple comparisons test). In addition, a wide literature relates late complex spikes occurring after movement initiation to movement errors and sculpting of simple spike rates during movement. Therefore, we analyzed kinematics in which we observed early or late complex spikes in these different epochs (see Methods). Trials with late Cspks had distinct kinematics compared to trials without late Cspks showing significantly deviated endpoints (Fig. 2b-e, N = 8 animals, n = 58 cells; Euclidean distance from session median, No Cspk trials vs. Cspk trials: p = 8.1 × 10^−4^, W = −847, r = −0.43, Wilcoxon signed rank test) but not peak velocity (Fig. 2f; p = 0.81, t = 0.24, d = 0.032, paired t-test). By contrast, reaches with early Cspks had no discernable kinematic differences (Fig. S4a,b; N = 8 animals, n = 58 cells; Euclidean distance from session median, No Cspk trials vs. Cspk trials: p = 0.32, W = −257, r = −0.13, Wilcoxon signed rank test; Peak velocity, No Cspk trials vs. Cspk trials: p = 0.75, t = 0.32, d = −0.042, paired t-test), although we cannot rule out changes in reaction time or movement initiation^54,55^.

**Figure 2.**
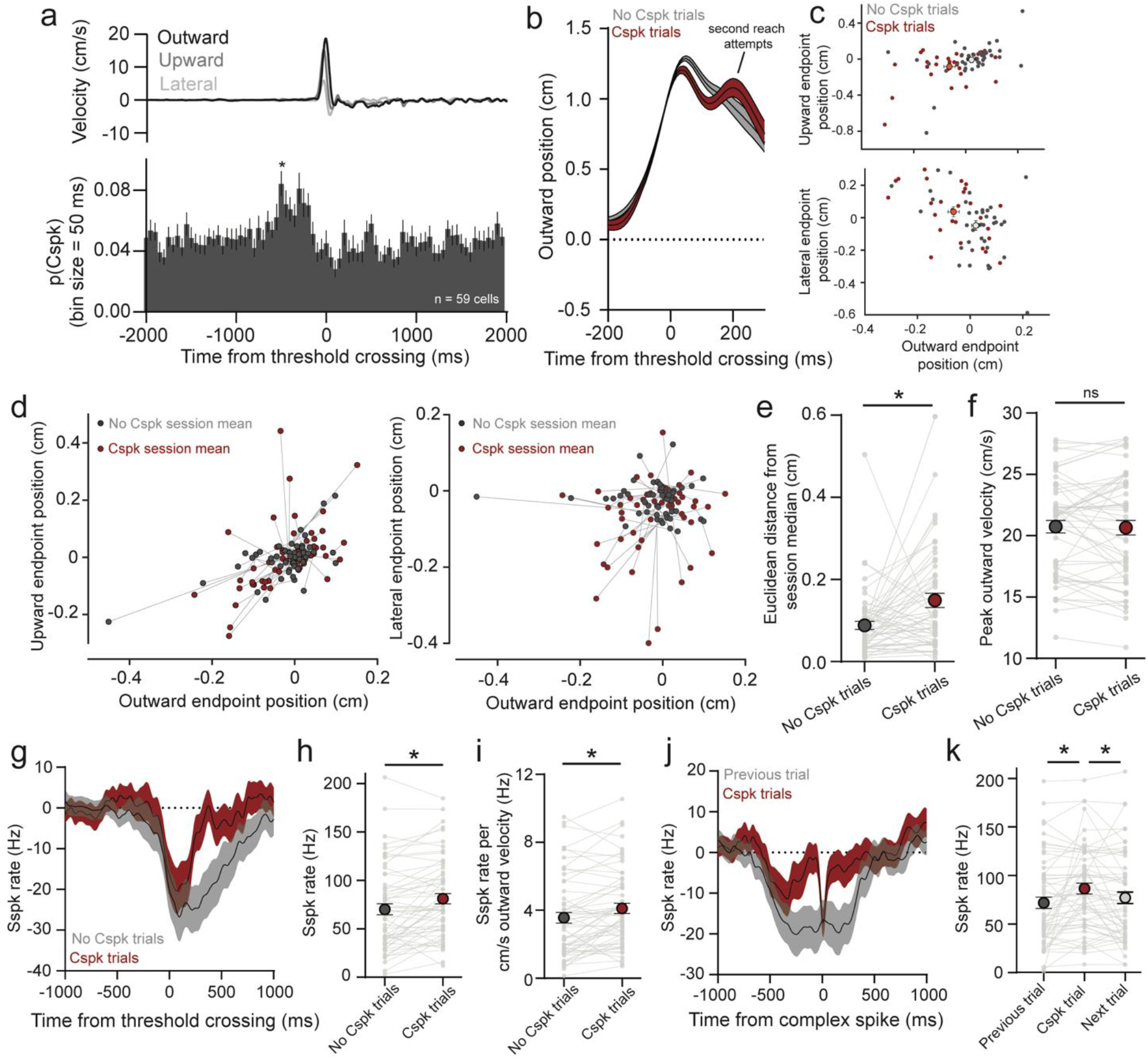
Reaches with Cspks have erroneous kinematics and elevated simple spike rates. **a.** Cspks are positively modulated in the 500 ms before reach before dropping close to baseline values. Top: Mean velocity of reaches with Cspks recorded. Bottom: PETH of Cspk activity relative to the time of threshold crossing (mean±SEM, asterisk indicates p < 0.05 for post-hoc Dunnett’s multiple comparisons test with mean Cspk firing rate). **b.** Positional profiles from an example session separated into reaches with (red) and without Cspks (black) during or shortly after outreach (mean±SEM). **c.** Endpoint of reaches relative to session median in the outward and upward directions (top) and outward and lateral directions (bottom) for trials with and without Cspks. Large red or grey dot indicates mean and SEM from Cspk and non-Cspk reaches. **d.** Session endpoints relative to session median for Cspk and non-Cspk reaches for each recorded cell with Cspks during or after outreach (n = 58 cells). Grey line links Cspk endpoint average with non-Cspk endpoint average for an individual session with the recorded cell. Left: outward and upward endpoint position. Right: outward and lateral endpoint position. **e.** Reach endpoints on Cspk trials were significantly further from session median compared to non-Cspk trials. **f.** Peak outward velocity was not significantly different between Cspk and non-Cspk trials. **g.** PC simple spikes (Sspk) on Cspk and non-Cspk trials aligned to threshold crossing (mean±SEM). **h.** PC simple spike (Sspk) rates were significantly higher during outreach in trials with Cspks (n = 58 cells). **i.** Ratio of simple spike rate to outward velocity was significantly higher during outreach in trials with Cspks. **j.** Simple spike rate aligned to the time a Cspks, or simple spikes aligned to the same time relative to threshold crossing on the previous trial showed simple spike increases shortly before the Cspk (mean±SEM). **k.** Quantification of simple spike rates in the 100 ms before a Cspk on a Cspk trial or a the previous/next trial aligned to the same time of the Cspk relative to threshold crossing.

To test whether the relationship of PC rates to kinematics changes on Cspk trials, indicative of an encoding error, we compared simple spike rates on Cspk and non-Cspk trials for early and late Cspks. Across neurons, simple spike rate was significantly elevated on late-Cspk trials (Fig. 2g,h; No Cspk trials vs. Cspk trials: p = 2.8 × 10^−4^, t = 3.9, d = −0.51, paired t-test) and this elevation led to a shift in the relationship of simple spike rate to reach velocity (Fig. 2i; No Cspk trials vs. Cspk trials: p = 1.0 × 10^−3^, W = −833, r = −0.42, Wilcoxon signed rank test). Trials with early Cspks did not show elevated simple spike rates during outreach or changes in the relationship between simple spikes and reach velocity across sessions (Fig. S4c-e; Simple spike rate during outreach, No Cspk trials vs. Cspk trials: p = 0.37, t = 0.90, d = −0.12, paired t-test; Simple spike rate to peak velocity ratio, No Cspk trials vs. Cspk trials: p = 0.29, W = −273, r = −0.14, Wilcoxon signed rank test). Cspks function to depress PC inputs, leading to reductions of simple spike rate^56,57^. If Cspks are responding to erroneous simple spike elevation, we speculated that simple spike rate should be elevated shortly before the time of a Cspk, as has been previously demonstrated^58^. We therefore analyzed simple spike rates aligned to the time of the Cspk, or the same time on the previous or next trial. In late Cspk trials, Cspks were associated with higher than average simple spike rates in the 100 ms before a Cspk compared to the previous trial, and simple spikes in this window were lowered on the trial after the Cspk trial (Fig. 2j,k; p = 9.8 × 10^−4^, F = 8.0, RM one-way ANOVA; previous trial vs Cspk trial: p = 3.6 × 10^−3^, d = −0.45, Cspk trial vs next trial: p = 0.024, d = 0.36, Tukey’s multiple comparisons test). In contrast, early Cspks that occurred before the onset of reach did not display increases in simple spikes before the Cspk (Fig. S4f,g; p = 0.17, F = 0.69, RM one-way ANOVA). Together these data reveal dynamics of PC Cspks, simple spikes, and associated kinematics that suggest a continuous recalibration of kinematic tuning in PCs.

### Repeated closed-loop optogenetic perturbation of cerebellar inputs during reach causes hallmark characteristics of sensorimotor adaptation

Next, we sought to probe whether PCs reweight cerebellar inputs that shape movement kinematics. Previous work has shown that stimulation of pontine afferents to the cerebellum perturbs reaching movements in mice^35^. This effect is interpretable as corrupted cortical information entering the cerebellum which initially drives an erroneous cerebellar control policy observable in acute kinematic effects. If cerebellar associative learning mechanisms implement the formation of an anticipatory control policy, a number of predictions emerge: pontocerebellar mossy fiber stimulation that drives reach errors will, when repeated over many reaches, lead to adaptation of PC responsivity. Removing the perturbation should lead to aftereffects due to accumulated learning of new contingencies. Finally, adaptation and aftereffects will be dependent upon the temporal context of the perturbation within the movement, where learning only accumulates when perturbations are temporally locked to the execution of the movement.

To drive erroneous activity in PCs during reaching movements, we injected AAV-expressing hSyn-ChR2 into the pontine nuclei in mice, a major hub relaying motor commands from motor cortex to the cerebellum^35,59–63^ (Fig. 3, Fig. S5a, Fig. S6a,b). Recordings of PCs showed that optogenetic stimulation of mossy fiber afferents in the cerebellar cortex drove both increases and decreases in simple spike firing rates (Fig. S5b; N = 4 animals, 43/151 cells, 26 increase, 17 decrease; p<0.05, paired t-test). This diverging stimulation effect is likely due to network properties in the cerebellar input layers leading to either net excitatory or inhibitory drive onto PCs^35,64^. Interestingly, cells with sorted Cspks (see Methods) showed a small but significant increase in Cspk probability in the 250 ms after stimulation during rest compared to the probability outside of this epoch in response to mossy fiber stimulation (Fig. S5c; N = 4 animals; n = 39 cells; p = 5.6 × 10^−3^, t = 2.9, d = −0.45, paired t-test), consistent with previous findings during electrical stimulation of mossy fibers^65^. Cspks time-locked to mossy fiber stimulation suggest that optogenetically-driven simple spikes m ay engage plasticity mechanisms to respond to perturbation.

**Figure 3.**
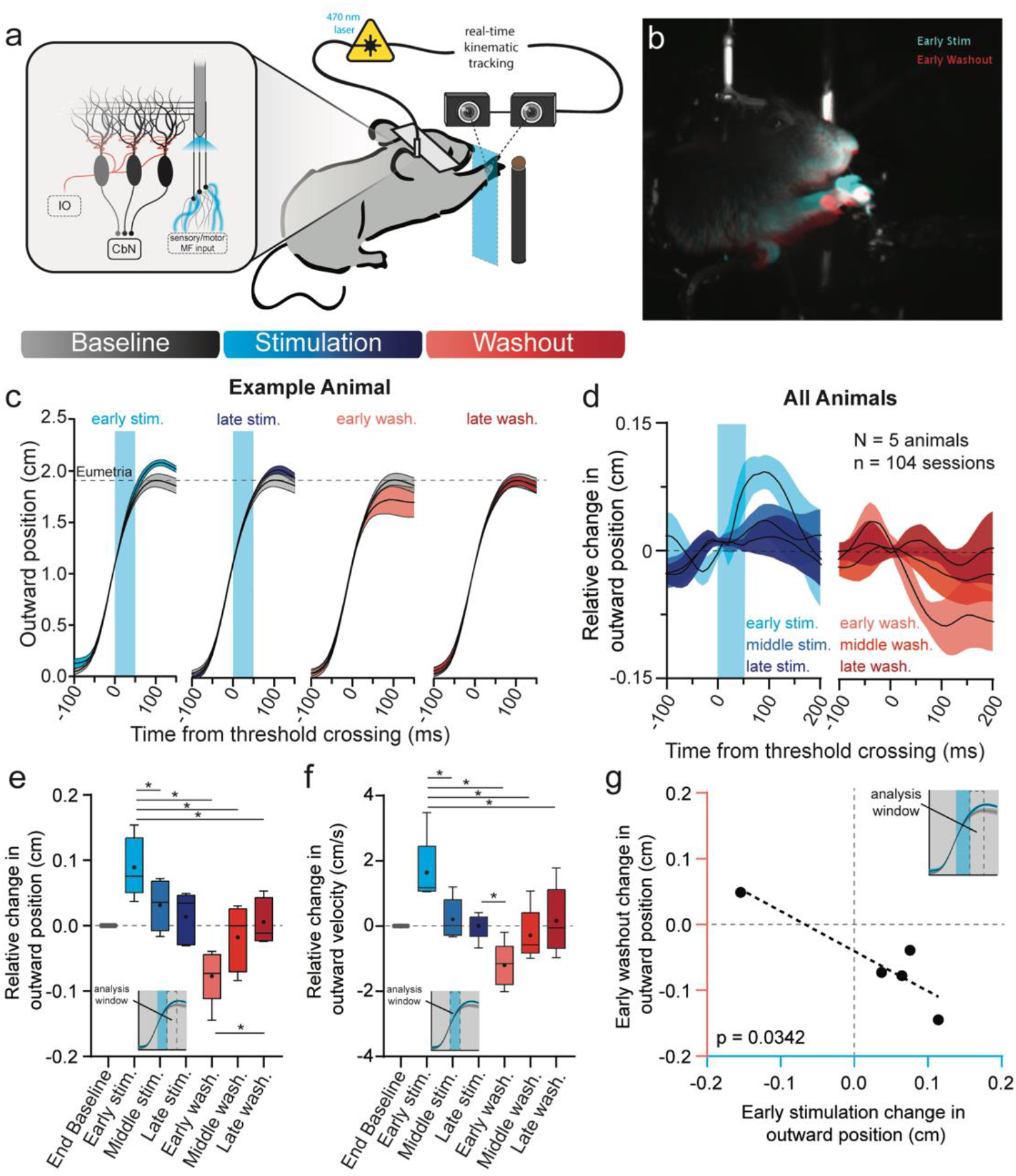
Adaptation to mossy fiber stimulation during reach. **a.** Headfixed mice expressing ChR2 in pontocerebellar mossy fibers were trained to reach for food pellets while the hand was tracked with high-speed cameras. On laser trials, light directed to cerebellar primary fissure through an implanted fiber was triggered in closed loop after the hand crossed a plane 1 cm outward from reach start position. Bottom: Perturbation schedule followed canonical adaptation structure, with a baseline (no-stimulation) block, stimulation block with stimulation on every reach, followed by a washout block with stimulation omitted. **b.** Hand position 100 ms after threshold crossing in the first stimulated (blue) and washout (red) reaches heading to the target (white), after Guo et al. (2021). **c.** Hand position during baseline (grey), compared to hand position measured across the adaptation and washout blocks in an example mouse (n = 20 sessions; mean±SEM). Blue shading denotes the time of mossy fiber stimulation. **d.** Summary of stimulation-induced kinematic effects, which decay over the adaptation block and show opposing aftereffects. Baseline subtracted hand position, rectified relative to the direction of kinematic effect of stimulation, is shown for reaches in the early (first reach), middle (middle 5), and late (last 5) phases for both stimulation (blue) and washout (red) blocks (N=5 mice; 104 sessions; mean±SEM). **e.** Summary of adaptation effects across animals and sessions. Relative change in outward position was assessed in the 50-ms window following the end of stimulation. Asterisks indicate statistically significant differences between blocks (p values reported in main text; Box and whiskers denote median, 25%, 75%, max and min, circle indicates mean). **f.** Same as **e**, but with outward velocity assessed in the 50-ms following the start of stimulation. **g.** The magnitude and direction of early stimulation effect was related to aftereffects. Plot shows linear regression relating the magnitude of the early stimulation outward position effect and early washout outward position effect compared to baseline reaches.

To assess whether repeated closed-loop stimulation could engage cerebellar learning mechanisms to produce sensorimotor adaptation, optical fibers were implanted in cerebellar cortex at the interface between Lobule Simplex and Lobules 4/5 (Fig. S6c,d). Experiments were structured in a block format where animals reached unperturbed in a baseline block, followed by a stimulation block where closed-loop stimulation of pontocerebellar axons (50-ms train) was delivered on every reach when the hand passed a 1-cm threshold in the outward direction, and finally a washout block where stimulation was removed to assess any aftereffects of learning. Each block was roughly 15-30 reaches long determined by each individual animal’s endurance in the task (Fig. 3a; baseline: 23.1 ± 6.24 reaches; stimulation: 22.4 ± 5.77 reaches; washout: 20.56 ± 6.65 reaches; mean ± SD; N = 5 animals, 104 sessions). Early in the stimulation block, we found that stimulation caused acute changes in reach kinematics: in 4/5 animals it caused hypermetric reaches in outward position and in 1 animal it caused hypometric reaches (Fig. 3b-c, Fig. S7a examples 1 and 2). To assess the relative change in hand position over the stimulation block, we measured the magnitude of the stimulation effect over the block, defining the initial direction of the stimulation effect on hand position as positive and the opposing direction as negative. We found that the magnitude of the stimulation effect decreased over the stimulation block. When the stimulation was removed, early in the washout block reaches deviated in the direction opposite the initial stimulation direction, before eventually correcting back to baseline at the end of the washout block in both outward position and velocity (Fig 3d,e; N = 5 animals, 104 sessions; Outward position: p = 1.7 × 10^−3^, F = 11.3, RM one-way ANOVA; early stimulation to middle stimulation: p = 8.9 × 10^−3^, d = 2.0; early stimulation to early washout: p = 0.030, d = 1.5; early stimulation to middle washout: p = 0.017, d = 1.7; early stimulation to late washout: p = 0.031, d = 1.4; early washout to late washout: p = 0.042, d = −1.3, Tukey’s multiple comparisons test; Outward velocity: p = 4.7 × 10^−3^, F = 12.1, RM one-way ANOVA; early stimulation to middle stimulation: p = 0.024, d = 1.6; early stimulation to early washout: p = 0.041, d = 1.3; early stimulation to middle washout: p = 3.5 × 10^−3^, d = 2.6; early stimulation to late washout: p = 0.030, d = 1.5; late stimulation to early washout: p = 0.048, d = 1.3, Tukey’s multiple comparisons test). The magnitude of the initial stimulation effect on outward position predicted the magnitude of the initial washout aftereffect across animals (Fig. 3f, R^2^ = 0.820, slope = −0.608 with 95% CI [−1.13, −0.0853], p = 0.034, F = 13.7); however, hypometric effects were generally larger than hypermetric effects (both during stimulation and washout), possibly due to biomechanical constraints of the limb and reaching apparatus imposing a ceiling effect on hypermetric movements. Interestingly, the aftereffect did not appear until the time that stimulation would have been delivered during outreach (Fig. 3c, Fig. S7d). In control experiments using red light (635 nm) we observed no kinematic deviations or adaptation profiles as seen with blue-light stimulation (Fig. S7e). Further, blue-light stimulation at rest produced negligible movements (Fig. S7f; N = 4 animals, 21 sessions; maximum outward velocity during stimulation: 0.26 cm/s).

To summarize, we have shown that animals adapt to a precisely timed internal perturbation of pontocerebellar mossy fibers and this learning is reflected in opposing aftereffects when the perturbation is removed. Adaptation was temporally precise, with changes in limb kinematics early in the washout block timed to the predicted point of perturbation.

### PC recordings show electrophysiological correlates of adaptation at the time of perturbation

To investigate cellular correlates of learning in PCs during behavioral adaptation to this circuit-level perturbation, we performed stimulation experiments while recording near the optical fiber with a Neuropixel probe (Fig. 4a). To assure that any firing rate changes were not attributable to unstable cell isolation across the experiment, we assessed the stability of every PC using two metrics: a correlation of spike template waveforms and the displacement of units along the electrode in the baseline and washout blocks (Fig. S8, see Methods). 203 of 314 putative PCs were stable across the experiment, 159 of which were modulated with reach. In the analyses that follow, we analyzed stimulus responsivity over the stimulus block in all stimulus-responsive PCs and population-level changes over adaptation of all reach-modulated PCs. First, to assess optogenetic stimulus responsivity in these neurons, we compared simple spike firing rates between baseline and stimulated reaches within the 50-ms stimulation epoch. Consistent with mossy fiber stimulation at rest, we observed a diverging effect pattern with stimulation during reach: 17 cells showed significant increases in simple spike firing and 25 cells showed decreases (Fig. 4b, p < 0.05, paired t-test). In both groups, the efficacy of stimulation dropped over the course of the stimulation block, consistent with adaptation (Fig. 4c,d). To statistically analyze the progression of the stimulus effect over the stimulation block, we defined the direction of the initial response as positive for all cells, (pooling cells that were inhibited and excited by stimulation), then measured the response magnitude over time. The response magnitude dropped across the stimulation block such that in later trials, firing rates were not significantly different from baseline (Fig 4e,f; n = 5 animals, 42 stimulation modulated cells; p = 7.2 × 10^−3^, F = 5.5, RM one-way ANOVA; end baseline vs. first 5 stim: p = 2.6 × 10^−3^, d = −0.56; end baseline vs. middle 5 stim: p = 0.037, d = −0.41; end baseline vs. last 5 stim: p = 0.32, d = −0.26, Tukey’s multiple comparisons test). Notably, stimulation affected cells did not show consistent aftereffects opposing the direction of the initial stimulation effect when the perturbation was removed.

**Figure 4.**
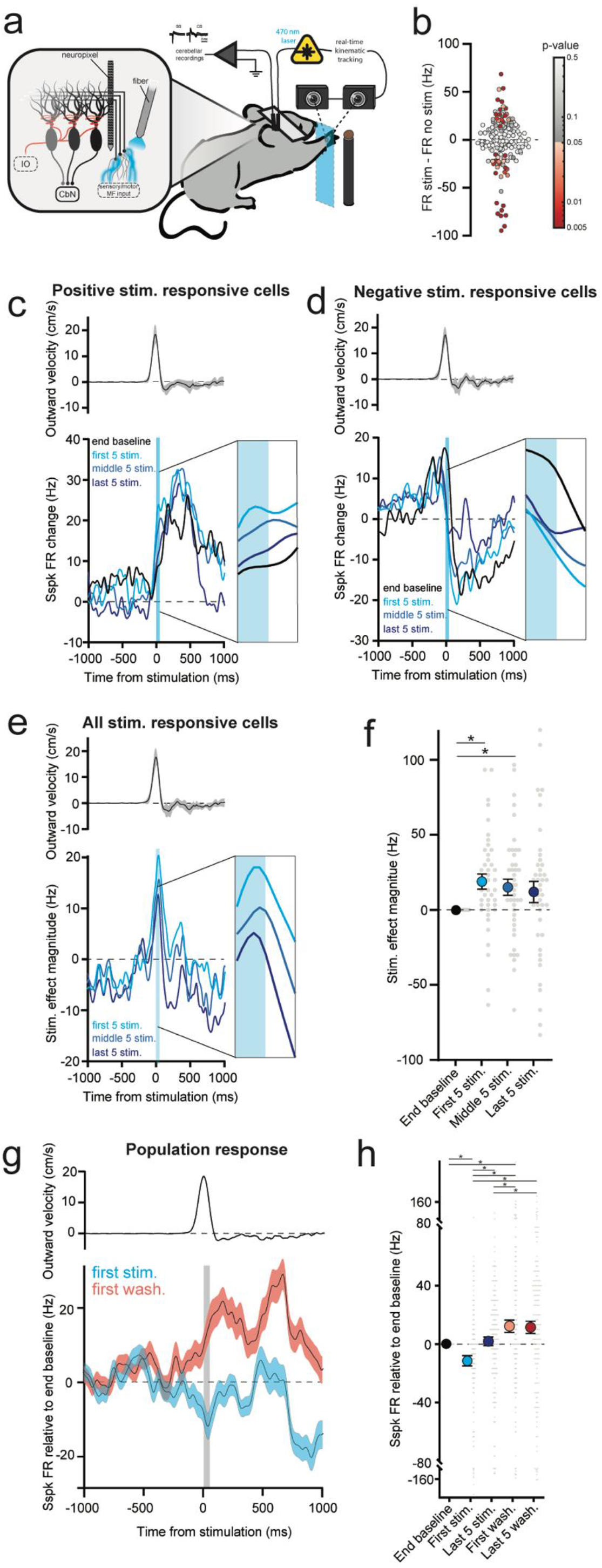
PCs show electrophysiological correlates of behavioral adaptation over the stimulation and washout blocks. **a.** Mossy fibers were stimulated at threshold crossing during outreach while recording PCs with Neuropixel probes. **b.** Mossy fiber stimulation effect during reach of all reach-modulated PCs. The difference in simple spike rate during the stimulation window is compared to the same epoch during baseline reaches (n = 159 cells). Significant differences are denoted by the color map on the right. **c.** Population summary of activity of PCs firing rate adaptation over stimulation block for all PCs positively modulated by stimulation. Top: mean reach velocity for all sessions (mean±SEM). Bottom: Average change in simple spike rates for the last 5 baseline reaches (black) and the first 5 (cyan), middle 5 (light blue), and last 5 (dark blue) stimulated reaches. n = 17 cells. **d.** Same as in **c** but for the population of PCs negatively modulated by stimulation. n = 25 cells. **e.** Same as in **c** but measuring the magnitude of stimulation across stim increase and stim decrease cells. Here the effect of stimulation is measured in the direction of the initial stimulation effect, thus a positive deflection for stimulation increase cells means an increase in firing rate relative to baseline, and a positive deflection for stimulation decrease cells means a decrease in firing rate relative to baseline. **f.** Quantification of the data shown in **e.** (mean±SEM). **g.** Population activity across all reach modulated cells. The first stimulated trial shows a negative deflection in net firing rate relative to baseline. Conversely, the first washout reach shows a net positive deflection. Grey box indicates the time of stimulation or analogous time in the washout block. **h.** Quantification of simple spike firing rates in the stimulation window for the data shown in **g** and the last 5 stimulated reaches and washout reaches.

Next, we analyzed how mossy fiber perturbations affected simple spike firing across the population of all reach modulated PCs (stimulus responsive and non-responsive cells). We observed transient effects of stimulation and opposing aftereffects that were visible on the first trial of the stimulation and washout blocks, respectively (Fig. 4g). Across the population, the net effect of the first stimulation was a reduction of simple spike firing rate relative to baseline. (Fig. 4g,h; n = 6 animals, 159 reach modulated cells; rates during the stimulation epoch: p = 2.3 × 10^−9^, F = 13.8, RM one-way ANOVA; end baseline vs. first stim: p = 0.012, d = 0.26; end baseline vs. first wash: p = 0.039, d = −0.23; first stim vs. last 5 stim: p = 7.4 × 10^−4^, d = −0.47; first stim vs. first wash: p = 3.0 × 10^−6^, d = −0.42; first stim vs. last 5 wash: p = 1.9 × 10^−7^, d = −0.47; last 5 stim vs. first wash: p = 0.019, d = −0.25; last 5 stim vs. last 5 wash: p = 0.015, d = −0.25, Tukey’s multiple comparisons test). This effect was rapidly adapted such that by the end of the stimulation block mean simple spike firing returned to baseline levels. On the first washout reach, there was marked increase in simple spike rates, an aftereffect opposite the direction of the initial stimulation effect. This aftereffect was only marginally lower by the end of the washout block; however simple spike firing outside of the stimulation window showed a more visible normalization to baseline levels (Fig. S9a). The dataset was underpowered to relate complex spike probability to these changes, but in the PCs in which complex spikes were observed, the mean Cspk rate in the 250 ms after stimulation was not significantly different across the blocks (Fig. S9b; N = 5 animals, n = 13 Cspk sorted cells, p = 0.31, F= 1.2, RM one-way ANOVA). Overall, these data demonstrate acute effects of stimulation that adapt across the stimulation block and population-level net aftereffects that oppose the initial firing rate deflection caused by stimulation, consistent with adaptation to perturbation and opposing aftereffects seen in reaches.

### Dissociation of adaptation and aftereffects with a randomized perturbation schedule

In the experiments above, we have shown that adaptation is temporally specific (e.g. Fig S7b). We hypothesized that the temporal specificity of perturbation within the reach produced a fixed association between active inputs and error, facilitating adaptation. We therefore predicted that by presenting spatially inconsistent stimuli trial to trial, mice would not adapt to stimulation. To test this, we repeated block-stimulation experiments, but rather than stimulating when the hand passed the 1-cm outward plane, we stimulated at a pseudorandomized position in the outward direction uniformly distributed between 0.3 and 1.8 cm (Fig 5a,b). To assess the effect of stimulation at different points in the reach, we aligned reaches to the time of stimulation and measured the difference in position compared to aligned baseline block reaches. Baseline subtracted reach profiles showed a characteristic change in outward position aligned to the time of stimulation, similar to results in fixed-position stimulation experiments. Surprisingly, even though perturbation positions were distributed across the stimulation block, we found that animals still exhibited adaptation to the stimulation early in the stimulation block, although this adaptation plateaued to intermediate levels between middle and late block epochs in outward position and velocity (Fig. 5c,d n = 5 animals, 60 sessions; Outward position: p = 0.016, F = 10.8, RM one-way ANOVA; baseline to early stimulation: p = 3.6 × 10^−4^, d = −3.9, Tukey’s multiple comparisons test; Outward velocity: p = 0.016, F = 7.5, RM one-way ANOVA; baseline to early stimulation: p = 0.040, d = −1.1; early stimulation to late stimulation: p = 0.017, d = 1.4, Tukey’s multiple comparisons test). To assess the presence of aftereffects, we analyzed the positional and velocity differences between baseline and washout reaches near the mean of the distribution of stimulus thresholds (50-100 ms after crossing the 1-cm outward plane). Despite evidence for adaption to the randomized stimulation, there were no consistent aftereffects; instead, reaches tended to have a greater distribution of positional differences that averaged to roughly zero (Fig. 5e, Fig. S10; Outward position: p = 0.65, F= 0.40, RM one-way ANOVA; Outward velocity: p = 0.76, F= 0.23, RM one-way ANOVA).

**Figure 5.**
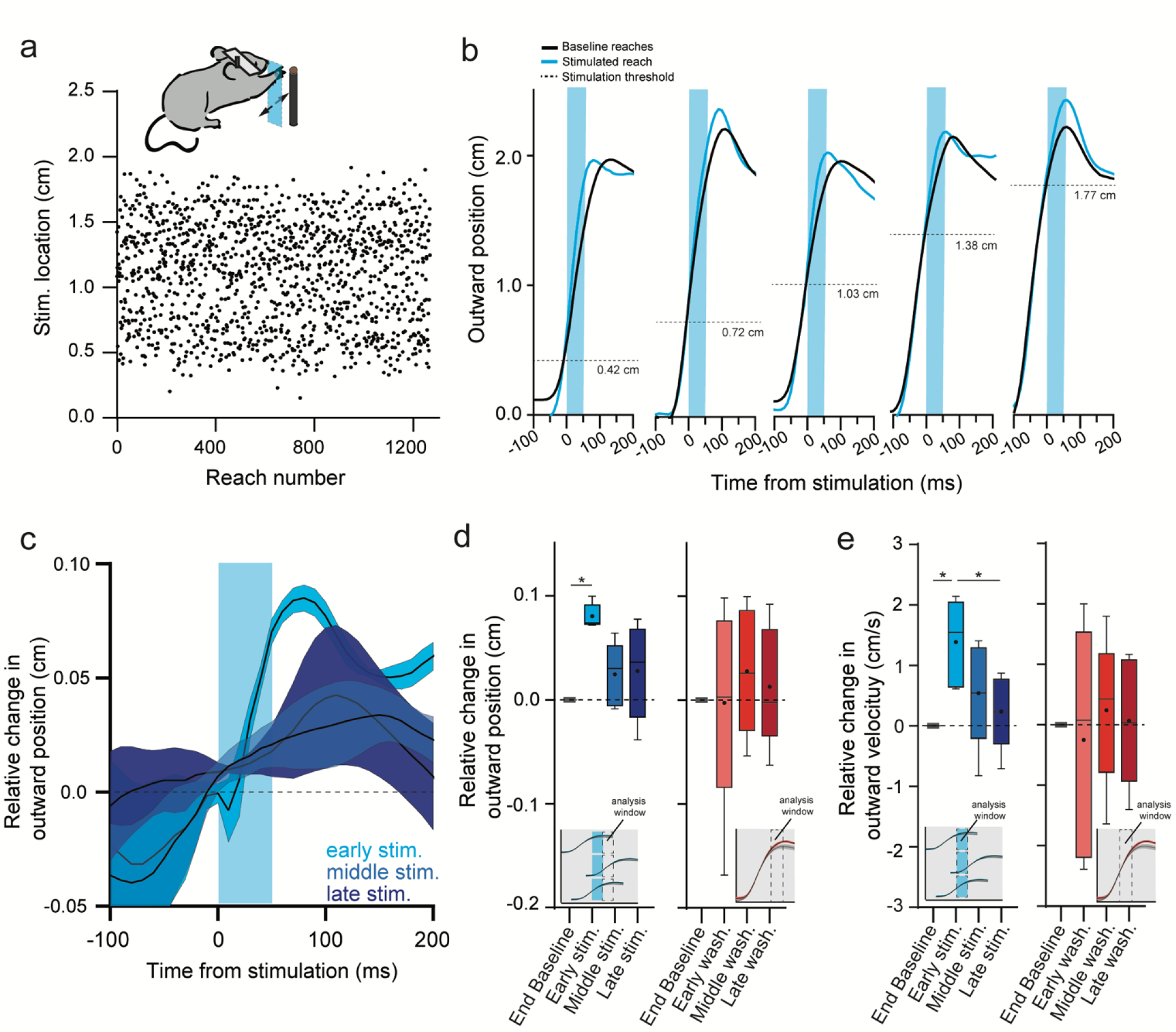
Dissociation of adaptation and aftereffects with randomized stimulation position. **a.** Stimulation location during outreach was distributed pseudorandomly between 0.3 and 1.8 cm in the outward direction during the stimulation block. **b.** Examples of reaches stimulated at 5 different locations during outreach. Each stimulated reach is compared to the last 5 baseline reaches of each session. The horizontal dashed line indicated the threshold crossing that triggered stimulation. **c.** Summary data of relative change in outward position for stimulation reaches in the early, middle, and late block. N=5 mice; 60 sessions. **d.** Quantification of stimulation effect on outward position across adaption block. For each reach, the analysis window was the 50-100 ms after stimulation onset aligned to the time of threshold crossing for each reach (inset). Quantification of aftereffects on outward position during washout block. Here, the analysis window is the 50-100 ms after crossing the 1-cm threshold for each reach – the same as the analysis in fixed-position stimulation experiments. **e.** Same as **d** but instead quantifying of outward velocity in the stimulus window, and aftereffects in the 50 ms after crossing the 1-cm threshold for each reach.

### A cerebellar model accounts for experimental adaptation and aftereffect dissociation

To better understand the non-intuitive adaptation profile of position-randomized stimulation, we modified a simple model of PC firing based on a previously published study^66^. The model takes as an input parallel fibers and inhibitory interneurons, each active for 15 ms, that as a population tile a 400-ms hypothetical movement (Fig. 6a). The PC rate mimicked the net firing rate suppression that we see in population activity during reach. At equilibrium, the populations of parallel fibers and interneurons are perfectly balanced during the movement and cause no deviation in the PC firing rate from trial to trial. The model employed a learning rule such that any elevation of the PC rate from this equilibrium would lead to depressing the weights of parallel fibers active at the time of deviation through a Cspk-like error signal, as in cerebellar LTD. Conversely, parallel fibers with depressed weights relax back to baseline levels in the absence of Cspks. We titrated the learning rate to match that observed in fixed-position stimulation experiments (see Methods).

**Figure 6.**
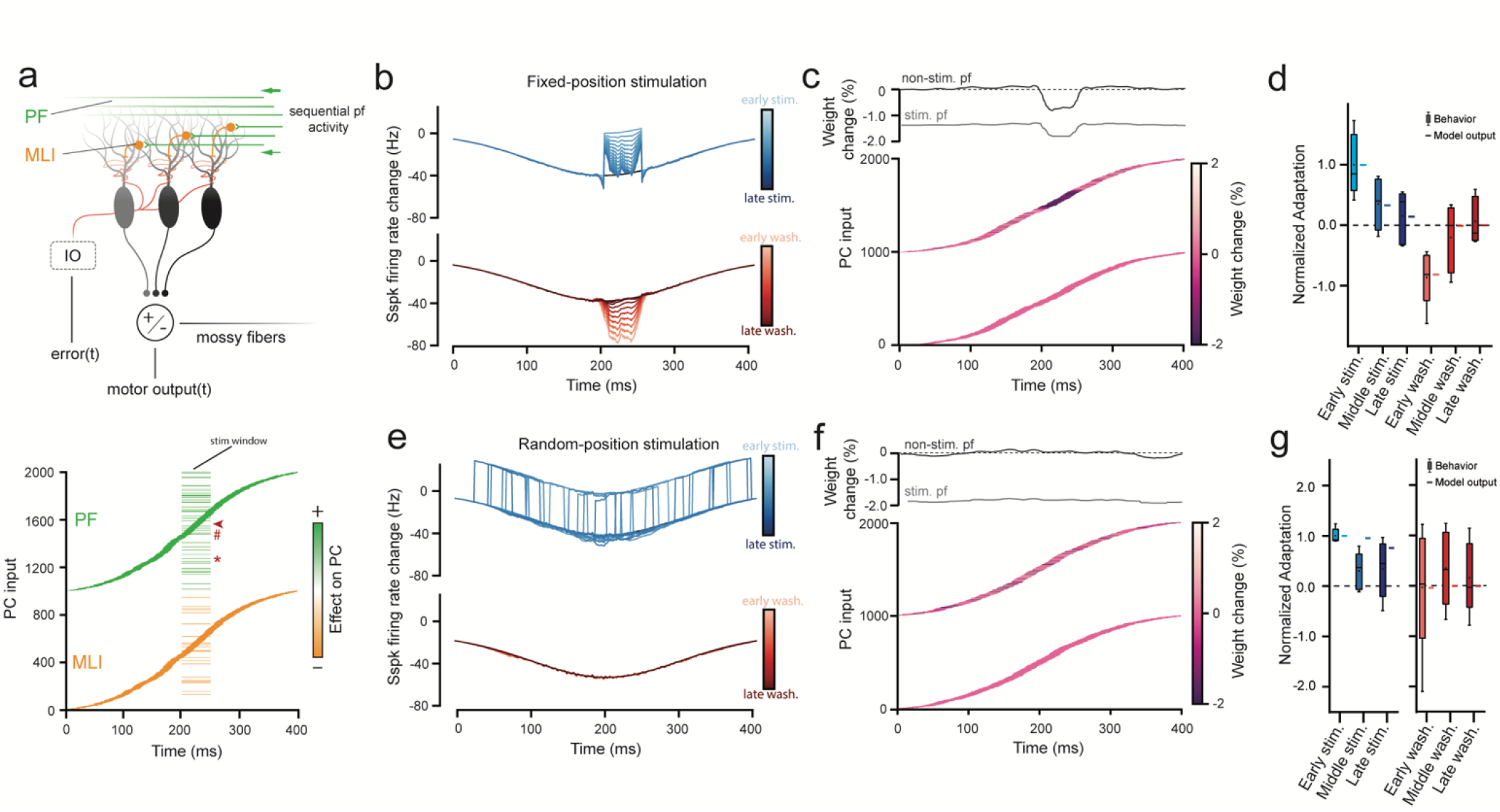
A cerebellar model accounts for adaptation and aftereffect dissociation. **a.** Schematic diagram of the temporal cerebellar-learning model. The model input is a population of 2000 cells, divided into 2 balanced populations of 1000 parallel fibers and 1000 interneurons, activated during a brief window during a simulated 400-ms movement. The output of the PC module that receives this information is compared to the input in the cerebellar nuclei. At equilibrium, the 2 populations are perfectly balanced (parallel fibers cause net activation of the PC, and the interneurons cause a net decrease; bottom) and the PC module outputs an activity curve (Gaussian that mimics the firing rate suppression observed in empirical data) that spans the movement. Positive deviations from this curve (errors) lead to mismatch in the nuclei and subsequent activation of the inferior olive, which reduces the weights of parallel fibers active shortly before the error. To simulate optogenetic perturbation experiments (bar-code like pattern at 200 ms), a step of activity was added to a subset of parallel fibers and interneurons for 50 ms in the center of the movement (fixed stim) or randomized across the block (random stim). Note that stimulation can either activate a cell twice (e.g. parallel fiber 1257, *) or overlap with endogenous activity (e.g. 1490, #), and non-stimulated neurons can be endogenously active during the stimulus window (e.g. 1561, arrow). **b.** PC simple spike activity during the stimulation block (top, blue) and washout block (bottom, red) showing progressively adapting response magnitudes during the adaptation block and progressively decaying aftereffects during washout. **c.** Parallel fiber weight changes at the end of the fixed-position stimulation block. Top: change in weights of “artificially” stimulated and non-stimulated parallel fibers plotted by time of endogenous activation. Bottom: heatmap of parallel fiber weight changes on top and unchanged interneurons on bottom. Note population weight change concentrated at time of stimulation, seen in both artificially stimulated and unstimulated fibers during stimulation epoch. **d.** Comparison of model output to empirical observations for fixed-position stimulus conditions (Fig.3). Model closely matches behavioral adaptation. **e.** Same as **b** but here the stimulation window is randomized across the reach. **f.** Same as **c** but for random position stimulation experiments. Note the absence of clustered weight changes in unstimulated parallel fibers. **g.** Comparison of model output to empirical observations for random-position stimulus conditions (Fig.5) showing that both model and empirical observations show adaptation but not directional aftereffects.

First, we modelled fixed-position optogenetic-perturbation experiments by artificially increasing activity in a random subset of parallel fibers and interneurons 50 ms in the middle of the hypothetical movement (Fig. 6a). Differential parallel fiber to interneuron activation ratios lead to a net activation of the PC to engage the Cspk-on learning rule, see Methods. Initially, this modification of PC inputs caused a large deviation in the PC firing rate in the stimulated window, resulting in an error and synaptic depression of the concomitantly active parallel fibers (Fig. 6b). Over several repeated perturbation trials this reweighting minimized the effect of the perturbation, correcting PC firing rate back to baseline. After 20 trials, we removed the perturbation. The model output then exhibited opposing aftereffects in PC firing rate at the previous time of perturbation, before eventually relaxing back to baseline. The adaptation profile was quantitatively similar to the empirically observed behavior. Importantly, we note that the aftereffect seen in the PC firing profile is a consequence of depressed weights in both perturbed parallel fibers and other unperturbed parallel fibers that were coincidentally active at the time of the perturbation (Fig. 6c,d). Thus, the model was unable to distinguish the difference between parallel fibers that caused or did not cause a deviation from the target PC activity within the perturbation epoch.

Next, we modeled the position-randomized mossy fiber stimulation paradigm (Fig 6e-g). As with the empirical results, we saw a reduction in the magnitude of the perturbation effect, consistent with high probabilities of Cspks around the time of a perturbation – that is, the perturbed inputs are subject to learning because they are always aligned to the error that follows (Fig. 6e). While the magnitude of adaptation was smaller than observed in the fixed position model, we found that the model learning plateaued late in the perturbation block, similar to empirical observations (Fig. 6g). When the perturbation was removed, there were minimal aftereffects, also consistent with experimental data. Model weights at the end of perturbation show that this absence of aftereffects is explained by the lack of accumulated learning in coincidentally active parallel fibers; *i.e.*, when perturbations are distributed across the movement, coincidently active parallel fibers are different from trial to trial, and therefore subjected to only transient plasticity (Fig. 6f). Thus, in randomized stimulation, the presence of adaptation illustrated a mechanism by which the cerebellum distinguishes cause-and-effect using time: adaptation is explained by the conserved causal relationship between stimulated PC inputs and error, while the absence of an aftereffect is the result of unaccumulated trial-over-trial learning in coincidentally active non-stimulated inputs. By contrast, aftereffects in the fixed position paradigm are a consequence of the system generalizing attribution of error to fibers that were merely coincidently active relative to perturbation but did not necessarily drive error.

## Discussion

Here we discovered a naturally occurring PC population suppression during mouse reaching movements that scaled with the velocity of outreach and occurred shortly before the transition to the decelerative phase of movement, reminiscent of emergent PC population kinematic coding in oculomotor vermis during saccades^45^. We speculate that this suppression is a type of conditioned response: sensorimotor information relayed through mossy fibers become learned cues for PCs to scale the decelerative phase of movement via disinhibition of the anterior interposed nucleus. We further demonstrate kinematic effects of mossy fiber stimulation that decrease over trials, akin to sensorimotor adaptation, with concordant changes in PC activity that imply cerebellar associative learning. We observed a surprising dissociation of adaptation and aftereffects when randomizing the position of stimulation during reach, designed to test the reliance of adaptation on perturbation context. A model demonstrated that aftereffects are a consequence of misattribution of error to consistently coactive parallel fibers. Conversely, the dissociation of adaptation and aftereffects reflects a lack of accumulated plasticity at a single point during the movement.

By demonstrating remapping of inputs to outputs of the cerebellar cortex, we link concepts developed in delay eyeblink conditioning to adaptation of a skilled volitional movement. Specifically, the mossy fiber stimulation used here to drive reach perturbations is analogous to mossy fiber stimulation used as a conditioned stimulus in eyeblink conditioning. We speculate that motor plan or early kinematic information acts endogenously as such a conditioned stimulus associated with reach outcome that, when erroneous, drives cerebellar learning^51^. We note some nuanced differences between paradigms, however. For instance, adaptation to pontocerebellar stimulation occurs within tens of trials, many fewer than conditioned eyeblink responses, which require hundreds of pairings ^67^. However, non-human primates and cats exhibit rapid adaptation consistent with our results in other sensorimotor adaptation paradigms^32,68^. One possible explanation for these different learning speeds is the richness of a temporal basis set that may emerge in the cerebellar granule cell layer in response to inflow of efferent commands and sensory feedback during movement compared to a relatively impoverished basis set from an invariant, unimodal conditioned stimulus. Indeed, locomotion concurrent with eyeblink training expedites learning^69^, consistent with this view. Further, eyeblink responses are generated *de novo,* where PCs must develop a novel response to an unfamiliar stimulus. Conversely, pontocerebellar stimulation during reach alters the execution of a movement for which a mossy fiber to cerebellar output mapping may already exist. Thus, PCs must simply adjust already existing responses, potentially speeding the rate of learning.

Another conspicuous departure from learning seen in eyeblink conditioning is that mossy fiber stimulation during reach drives an error. Thus, the unconditioned stimulus is not externally imposed but is rather the erroneous behavior that results from the perturbed mossy fiber activity. In this sense, the mossy fiber activity that interferes with cerebellar control acts as both a conditioned stimulus and generates a movement error that acts as an unconditioned stimulus to drive learning. In addition, eyeblink conditioning involves the cerebellum associating two stimuli that are not causally linked (*i.e.*, the tone does not cause an air puff) while reach adaptation associates sensorimotor information that is causal to reach error (*i.e.*, the erroneous motor commands generated by the cerebellum cause a reach error). Because this conditioned stimulus cannot be decoupled from the movement error, adaptation should always occur with repeated stimulation even under randomized stimulus conditions. In the case of external perturbations of limb movements, randomizing the direction of perturbations on reaching movements manifests as reach adaptation on the subsequent trial^70^, but adaptation does not accumulate because the cause of errors cannot be predicted. Our randomization experiments have a key difference: the internally perturbed mossy fibers are a consistent source of error, allowing the system to drive adaptation to these inputs. Importantly, because these perturbed inputs have no temporal correlation with the movement, no aftereffects are produced in the native population of mossy fibers active in the absence of stimulation.

Isolating a locus of skilled reach adaptation to the cerebellum poses an important conceptual hurdle. Cerebral cortex is a major input to the pontine nuclei – the focus of perturbation in this study – thus learning in motor cortex must be accounted for in cerebellar contributions to movement. Likewise, cerebellar outputs relay information back to motor cortex indirectly via thalamus^71–74^. Previous work has demonstrated that reach-associated pontocerebellar stimulation drives activity in motor cortex^35^, meaning each learner in this loop stays apprised of the activity in the other. Could plasticity sites outside the cerebellum account for our observations? Our data argue for a locus of learning in the cerebellum in two major ways: First, we observe a reduced efficacy of mossy fiber drive onto Purkinje cells over many repeated trials. A parsimonious explanation is that highly-plastic parallel fiber synaptic weights are changing during adaptation rather than cortical commands overriding these proximal perturbations. Second, if PC firing rate changes were caused by modulated afferents to the cerebellum, it would be difficult to reconcile such a mechanism with adaptation to randomized stimulation because these compensatory cerebellar inputs could not predict the time of stimulation.

How might multiple connected brain regions, all of which are implicated in learning, accomplish learning a task in parallel? In our study, mice were expertly trained when we introduced optogenetic perturbation of inputs. Thus, stimulating pontocerebellar fibers, we corrupted the relationship of action directed by motor cortex and the established cerebellar response tuned to that action. Through adaptation, the cerebellum learned to assist movements with these newly modified inputs as evidenced by the diminishing kinematic effect on the limb; when stimulation was removed, the novel mismatch of cortical and adapted cerebellar contribution to the movement again manifests as movement errors. Our data unite two threads of cerebellar theory, classical conditioning and motor adaptation under the umbrella of associative learning, where active inputs to the cerebellum can be flexibly reformatted to more accurately accomplish a goal.

## Methods

### Animals

All procedures followed National Institutes of Health Guidelines and were approved by the Institutional Animal Care and Use Committee at the University of Colorado Anschutz Medical Campus. Animals were housed in an environmentally controlled room, kept on a 12 h light/dark cycle and had ad libitum access to food and water except during behavioral training and testing as described below. Adult C57BL/6 (Charles River Laboratories) mice of either sex (11 females, 8 males) were used in all experiments.

### Surgical procedures

All surgical procedures were conducted under Ketamine/Xylazine anesthesia. After induction of anesthesia, the surgical site was cleaned and subcutaneously injected with bupivacaine (2.5 mg/mL). Pressure injections of approximately 150 nL of AAV2-hSyn-ChR2-mCherry were stereotaxically targeted to the left pontine nuclei (−4.0 mm anterior-posterior, −0.5 mm medial-lateral, −5.4 mm dorsal-ventral, measured from bregma) and animals were allowed to recover for a minimum of 8 weeks to ensure expression in mossy fiber terminals in the cerebellar cortex. Custom made aluminum head plates were affixed to the skull centered on bregma with luting (3M) and dental acrylic (Teet’s cold cure). Optical fibers (105 micron core diameter, Thor Labs) attached to a ceramic ferrule (1.25 mm, Thor Labs) were implanted into the primary fissure, between Lob 4/5 and Simplex (−6.25 mm anterior-posterior, 1.9 mm medial-lateral, measured from bregma) at a depth of 1.2 mm^13^. For recording experiments, a craniotomy was made medial to the fiber placement and a recording chamber was secured with dental acrylic as previously described^75^.

### Behavioral task

Animals were allowed a minimum of 2 days of recovery after head fixation surgery, then were food restricted to 80-90% of their baseline weight for reach training. Mice were habituated to the headfixed apparatus by presenting food pellets (20 mg, BioServ #F0163) that could be retrieved with their tongue, then pellets were progressively moved further from the mouth until animals began reaching for food. Pellets were positioned to the right of the animal to encourage reaching with the right forelimb and moved to a consistent position specific to each mouse ~1.2 – 2.5 cm from reach start. Sessions lasted until animals successfully retrieved 20 pellets or until 30 minutes had elapsed, whichever came first. Mice were trained for a minimum of 15 days and were considered fully trained once they could successfully retrieve 50% of pellets 3 days in a row.

### Kinematic tracking and closed-loop optogenetic stimulation

Hand position was tracked in real time using an infrared-based machine-vision motion-capture system (6 Optitrack Slim3U Cameras mounted with LED ring arrays, Motive software) at 120 frames-per-second as previously described^2^. Cameras were positioned in front and to the right of the animal and focused on the approximately 8 cm^3^ spatial volume that covered the reach area of the right forelimb. 1-mm diameter retroreflective markers were used for camera calibration and affixed to the mouse hand for kinematic tracking. A custom-built calibration wand and ground plane were used to set position and orientation of the cameras in Optitrack Motive software. Camera calibration was refined monthly to account for any drift of the cameras over time. Calibrations that reported a mean triangulation error <0.05 mm were considered passes. The spatial origin was set to be at the center of the bar where mice placed their hand during rest. Spatial blocking and camera detection thresholds were adjusted to prevent erroneous tracking of minimally infrared-reflective objects.

Real-time hand positions were streamed into MATLAB (2018a) with a latency under 1 ms. Custom-written MATLAB code was used to detect when the hand passed a positional threshold 1-cm outward from the bar where the mice rested their hand then send a ‘go’ signal to an Arduino microcontroller (Uno) which triggered a laser with TTL pulses. To ensure low-latency closed-loop stimulation we used an open-source C++ dynamic link library^76^ edited to reflect the parameters of laser stimulation (50-ms stimulation, 100 Hz, 2-ms train). This system has a closed-loop latency of 9.5 ms from the time of threshold crossing (120 fps camera frame rate, 0.5 ± 0.1 ms (mean ± SD) MATLAB-Arduino communication). Hand positions and stimulation times were streamed into MATLAB and saved for post-processing.

### Kinematic analysis

All kinematic analysis was performed using custom-written MATLAB code. First, erroneously tracked objects were removed using a nearest neighbor analysis, which assessed the closest markers in subsequent frames and removed others, to produce a single positional trajectory of the hand marker over time. Any dropped frames where the marker was not detected were interpolated over, then data were filtered using a 2^nd^-order low-pass Butterworth filter (10 Hz)^77^ using MATLAB’s zero-phase filter function “filtfilt”. Last, interpolated points were removed such that the filtered marker positional data reflected only data captured during the experimental session.

To segment continuous data into reaches, we found instances of the marker passing the 1-cm positional threshold in the outward direction and clipped 10-s segments centered on this time point. We defined outreach as the segment of this data from the time before threshold crossing that the hand exceeded 2 cm/s in outward and upward velocity to the time after threshold crossing where the hand stopped moving in the positive outward direction (outward velocity < 0 cm/s). Occasionally, the marker would become obscured behind the pellet holder during reach or spurious detection of the nose would jump the marker position to the nose and be detected as a reach. Therefore, to prevent against analyzing reaches that had large segments of data missing, any threshold crossings where the marker dropped greater than 25% of points between the start and end of outreach were not considered for further analysis.

Reach velocity and acceleration were calculated using the numerical gradient between position timepoints in each dimension. To produce aligned reach position curves, we interpolated data at 10 ms centered on the time the hand passed the 1 cm positional threshold crossing in outward direction. The effect of stimulation was assessed by measuring changes in stimulation and washout reaches (early, middle, and late) relative to the last 5 baseline reaches in the 50-ms interval following the end of stimulation. To assess the unadapted effect of stimulation or washout, early reaches were defined as the first reach in each block; middle and late reaches were the middle 5 and last 5 reaches of reach block, respectively. To align random-stimulation position reaches, we found the positional threshold of stimulation on each reach, then aligned stimulation reaches and baseline reaches to the time they crossed this boundary during outreach, averaged across reaches, then measured the difference in these curves, yielding the stimulation-aligned positional difference between end baseline and stimulation reaches. For washout reaches in random-position stimulation experiments, reaches were aligned to the time of the threshold crossing at 1 cm such that the aftereffect could be compared to fixed-position stimulation experiments. To account for varying effects of stimulation seen across animals (hypermetric and hypometric movements), the direction of positional change in early stimulation reaches relative to baseline for each animal in random- or fixed-position stimulation experiments was defined as the positive direction and the opposing direction as negative for that animal in each paradigm, allowing us to group data across animals with diverging effects. To assess the time course of stimulation effects within individual animals, we measured differences in position at each timepoint between the early stimulation reaches and baseline reaches using a Wilcoxon signed rank test.

### Electrophysiology recording procedure

Craniotomies were made over the cerebellum ipsilateral to the reaching arm of in fully trained animals. A custom-made recording chamber was implanted over the craniotomy, the brain was covered with triple-antibiotic cream (Globe), and the recording chamber was sealed with Quik-sil silicone (World Precision Instruments) such that it could be preserved for multiple recordings.

### Single electrode recordings

Single-electrode recordings were performed with 3-5 MOhm platinum/tungsten optrodes (Thomas Recording). Once animals were headfixed, the electrode was targeted to −6.25 mm anterior-posterior, 1.9 mm medial-lateral (measured from bregma) then lowered into the brain up to a depth of 1.8 mm with a motorized micromanipulator (Newport Motion Controller, Model 861). Signals were band-pass filtered at 300-5000 Hz, amplified with a MultiClamp 700A amplifier (Axon Instruments), then digitized (CED Power3 1401) and recorded with Spike2 software (CED). Once a putative PC was isolated, the brain tissue was allowed to relax for 15 minutes. Cell sorting was performed offline using Psort^78^.

### Neuropixel recordings

Neuropixels were lowered into the brain using a motorized micromanipulator (Sensapex uMp micromanipulator). Once the electrode shank spanned the putative PC layer, the tissue was allowed to relax for 15 minutes. Electrophysiology data was acquired using an OpenEphys system (https://open-ephys.org/gui). Data were sorted offline in Kilosort2^79^ and manually curated in phy (https://github.com/cortex-lab/phy).

### Neural data analysis

Following sorting, isolated units were analyzed offline using custom-written MATLAB code. In well-isolated single-electrode units, simple spikes and identifiable Cspks were sorted using Psort. To identify Cspks in Neuropixel recordings, we cross-correlated cells with high firing rates in the cortex with adjacent low-firing-rate clusters and looked for the presence of a Cspk-aligned simple spike pause and characteristic simple spike and Cspk waveforms. In many cells Cspks could not be identified across the length of the experiment. In these cases, we identified PCs based on cortical location and electrophysiological criteria using the firing rate, CV2, and median absolute difference from the median interspike interval (MAD)^80^. Cerebellar cortical cells with a firing rate > 40 spikes/s, CV2 > 0.20, MAD < 0.008 were labeled as PCs (Fig. S2). Using these metrics we were able to positively identify 94.9% of Cspk identified cells. We visualized these metrics in a 2-dimensional space using the tSNE function in MATLAB (parameters: distance = ‘euclidean’, exaggeration = 4, perplexity = 30, learning rate = 5000). Instantaneous firing rates for PCs were calculated by taking the inverse of the ISI between spikes, convolving with a 20-ms gaussian, then sampling at 10-ms intervals. In Neuropixel recording adaptation experiments we analyzed reach modulated PCs, defined as exhibiting a firing rate change during the reach epoch ≥ 1 standard deviation of the mean firing rate of the cell. Cell recordings from the baseline (unstimulated) block from cerebellar stimulation experiments during reach were included in the datasets in Figures 1 and 2. For analysis of pooled population firing rate data in Figure 1, we normalized reaches by velocity for each session, and binned them into velocity quintiles. Thus, each cell was equally represented across all velocity quintiles. To find the magnitude of the firing rate decrease in grouped population PC data, we found the minimum value of the population firing rate traces for each percentile bin within the peri-reach window (−500 ms to +500 from threshold crossing). We found the time of firing rate suppression by measuring the point at which each trace decreased firing by 50% from peak to nadir in this peri-reach window. We characterized early Cspks as those that occurred within 500 ms before reach onset, corresponding to roughly the time of Cspk elevation seen across cells (Fig. 2a). Late Cspks were characterized as those that occurred during outreach or the 500 ms window after the end of outreach.

In pontocerebellar stimulation experiments, to assure that observed simple spike adaptation was not the result of changing unit isolation across the experiment, we assessed unit stability with two metrics: waveform correlation and unit displacement across the experiment^81^. To assess waveform correlation, we isolated the template waveforms for each unit on the electrode with the greatest spike amplitude and the 32 surrounding electrodes (33 total). We averaged 1000 randomly selected spike waveforms for each channel from the baseline block and the washout block, concatenated waveform templates across the 33 channels, then correlated the concatenated waveforms from the baseline and washout blocks (Pearson correlation). As a shuffled control, we correlated concatenated templates from neighboring units in the baseline and washout block. Neighboring units were defined as those whose 32 surrounding electrodes overlapped with the unit of interest. PCs whose across experiment waveform correlation did not exceed the 99^th^ percentile (0.76) of the across-unit shuffled control correlation were excluded from further analysis.

To assess cell displacement across the experiment we calculated the position of unit (*x, y*) using

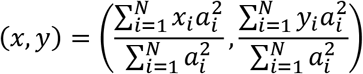

where *N* is the number of electrodes, *x*_*i*_ and *y*_*i*_ are the lateral and upward position of the electrode, and *a*_*i*_ is the peak-to-peak spike waveform amplitude on the *i^th^* electrode. Unit displacement was defined as the Euclidean distance between unit positions in the baseline and washout blocks. As a shuffled control, the displacement between neighboring units (as defined above) across the experiment was calculated. PCs whose displacement was above the 1^st^ percentile (2.36 μm) of shuffled control were excluded from further analysis.

### LASSO regression

To quantify the variance of PC simple spike firing rate that could be explained by reach kinematics, we used least absolute shrinkage and selection operator (LASSO) regression^36^. LASSO has the advantage of performing both regressor selection and regularization, producing a sparse model of many correlated kinematic regressors. 23 kinematic variables were used as regressors, including position, velocity, and acceleration in the upward, outward, and lateral directions, speed, unsigned acceleration, with each velocity and acceleration term additionally broken into positive and negative components. A full list of regressors is included in Fig. S3. Data for each reach were clipped into 2-s segments centered at the time of a 1-cm threshold crossing in the outward direction. Regression was performed using a custom-written MATLAB code using the “lasso” function. All kinematic data were standardized to have a mean of 0 and a variance of 1, and regression was performed with a 10-fold cross validation to avoid overfitting. To find the appropriate offset of firing rate and kinematics, instantaneous simple spike firing rates for each reach were offset by lags from 0 ms to −300 ms (firing rate leading kinematics) in 10-ms steps. The lag that minimized the mean squared error (MSE) of the regression was selected for each cell. To calculate the variance of firing rate explained, the predicted firing rates from the best fit regression were calculated from the kinematic data and compared to empirical data. R^2^ was calculated using:

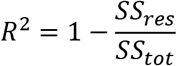

where *SS*_*res*_ is the sum of squared residuals and *SS*_*tot*_ is the total sum of squares.

For the spike shuffled control, spike times on individual trials were shuffled in time so that each reach epoch had the same mean firing rate, then converted to instantaneous firing rates as described above. For the reach shuffled control, reaches were assigned to firing rates recorded on different reaches. For both controls regressions were performed at the lag that minimized the MSE for empirical data and repeated 100 times; R^2^ values of each shuffled control were taken as the average of these 100 regressions. To assess the unique contribution of individual kinematic regressors to the fraction of variance explained in the empirical data regression, each regressor was time shuffled independently and regressions were repeated. The change in R^2^ value between the regressor shuffled regression compared to the complete empirical data model is the fraction of unique contribution to total variance explained for each kinematic variable^42^.

### Cerebellar model

The cerebellar model in the paper was derived from a previously published model^66^ and written using custom code in Python. A major difference between our paper’s model and the cited model is the assumption of a continuous temporal input of parallel fiber activity distributed across a hypothetical 400-ms movement, rather than a single parallel fiber input trial over trial. The model PC was fed 1000 parallel fibers that positively modulated the PC firing rate and 1000 MLIs that negatively modulated the PC firing rate, which were each active for 15 ms during a 400-ms interval, mimicking hypothesized temporal basis sets produced by the granule cell layer^82–84^. In the absence of perturbation these populations were perfectly balanced leading to no changes of PC firing from trial to trial. PC firing at time t on the n^th^ trial was calculated as the sum of the weighted contribution of all parallel fibers and MLIs at time *t*:

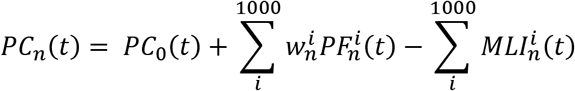

Here, 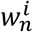 is the weight of parallel fiber *i* on the *n*^th^ trial and *PC*_*0*_ is the baseline firing rate of the PC.

Parallel fibers weights were subject to a learning rule based on deviations of the PC firing rate from trial to trial. Weights were adjusted following each trial according to two parameters: the probability of a Cspk as a function of trial error *βP*(*CS*|*E*_*n*_) where *β* (0.15) dictates the strength of synaptic depression in response to a Cspk, and a decay term, *α*_*PF*_ (0.95), that relaxes parallel fiber weights back to their initial value 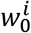:

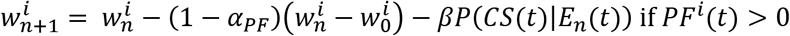

The probability of a Cspk is a function of *t*, where positive deviations in the PC rate from baseline at time *t* lead to elevation of Cspks rates from baseline leading to LTD and negative deviations of PC rate lead to reduction of Cspk rates from baseline levels, leading to LTP^85^. Specifically, the error at time *t* (*E*_*n*_ (*t*)) was used to calculate the probability of a Cspk at each time in the movement interval:

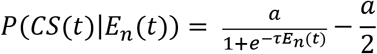

To obtain values for the parameters *a* and *τ*, we fit a curve to the change in position of early, middle, and late stimulated reaches in fixed-position stimulation experiments, then took the derivative of this curve to obtain the error correction (trial-over-trial positional change) for a given error magnitude.

We ran the simulations mimicking the experimental block structure used for empirical data, including a baseline block with no perturbation, an experimental block with a perturbation on every trial, and a washout block with the perturbation removed. For net positive perturbation trials, we added activity to a random subset of 150 parallel fibers and 50 MLIs at t = 200 ms for 50 ms that, when combined, drove an increase of 60 simple spikes/second in PCs at their initial weights (Fig 7). For net negative perturbation trials, we added activity to a random subset of 50 parallel fibers and 150 MLIs at t = 200 ms for 50 ms that drove a net decrease 60 simple spikes/second in PCs (Fig. S9). For each simulation, after 20 perturbation trials, the perturbation was removed, and the model was run for an additional 20 washout trials. To simulate random-position perturbation experiments, the time of perturbation was changed on every trial.

### Statistical analysis and data presentation

Data reported in the manuscript reflect statistical summaries from each animal across multiple sessions. For electrophysiological data, each neuron was treated as an independent sample. All data were tested for normality with the Kolmogorov-Smirnov test to choose the appropriate statistical analysis. All t-tests mentioned in the manuscript were two-sided, unless stated otherwise. In box and whisker plots, the box displays the median and 25^th^ and 75^th^ percentiles and the whiskers extend to the max and min of the data, with the mean is displayed as a dot in the box.

Effect sizes for parametric tests were estimated using Cohen’s d. For datasets with fewer than 50 samples, the Cohen’s d value was corrected for small sample size by multiplying by

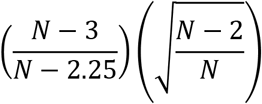

where N is the number of samples. Effect sizes for non-parametric tests were estimated by calculating r defined as

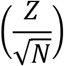

where Z is the Z statistic and N is the number of samples.

### Code and data availability

The code for cerebellar model can be found at github.com/dycala. Data used to make each of the figures is included in the supplementary materials. Analysis code and other data are available upon reasonable request to the authors.

## Acknowledgements

We thank the members of the Person lab for critical feedback on the manuscript; the Neurotechnology Center at the University of Colorado Anschutz Medical campus for use of core facilities including the Advanced Light Microscopy Core and the Optogenetics and Neural Engineering Core. We thank Mr. Michael Spindle and Ms. Elena Judd for technical assistance. Work was supported by F31 NS113395-01 to DJC; and NS114430, NSF CAREER 1749568, and by a grant from the Simons Foundation as part of the Simons-Emory International Consortium on Motor Control to ALP.

## Contributions

DJC and ALP designed experiments, interpreted data, and wrote the manuscript. DJC performed experiments, built the computational model, made figures, and analyzed data. MIB developed the closed-loop system, edited the manuscript, and contributed to early conceptualization of the project.

---

**Figure S1.**
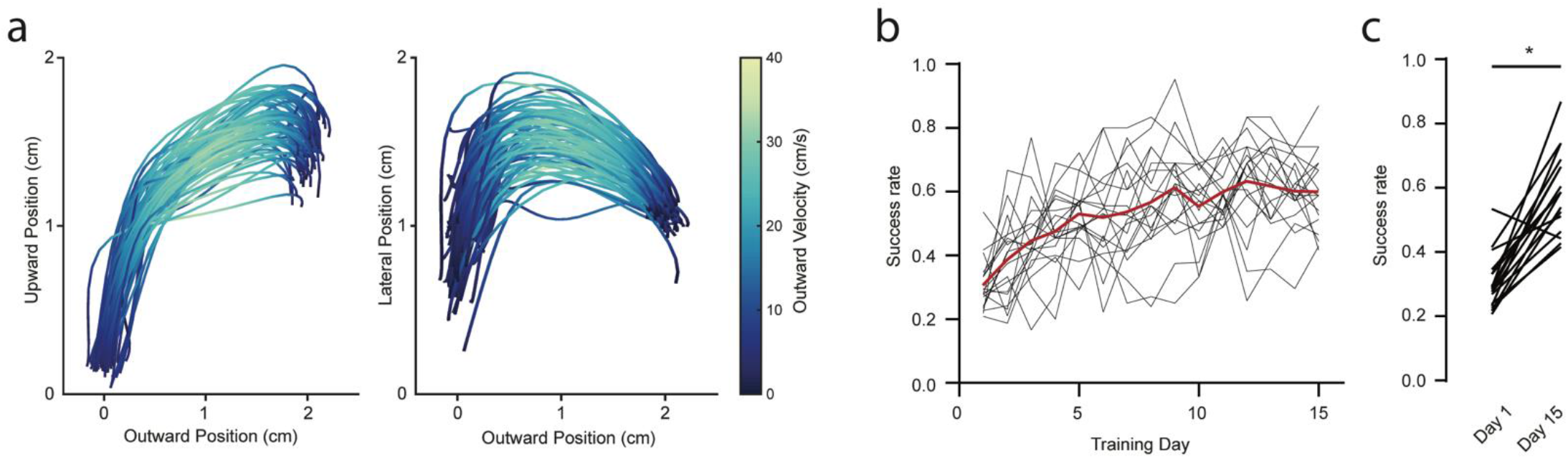
Reach tracking and reach performance over sessions. a. The right hand was tracked with high-speed cameras as mice reached upwards and outwards towards a food pellet. Positional outreach trajectories from a single session viewed are shown from a lateral (left) or bottom-up (right) vantage point with traces colored by the magnitude of outward velocity. b. Mice were trained for a minimum of 15 days on the reaching task. Pellet retrieval success was tracked throughout training for each mouse, mean is shown in red (n = 19 mice). c. Quantification of success rate on day 1 of training and day 15.

**Figure S2.**
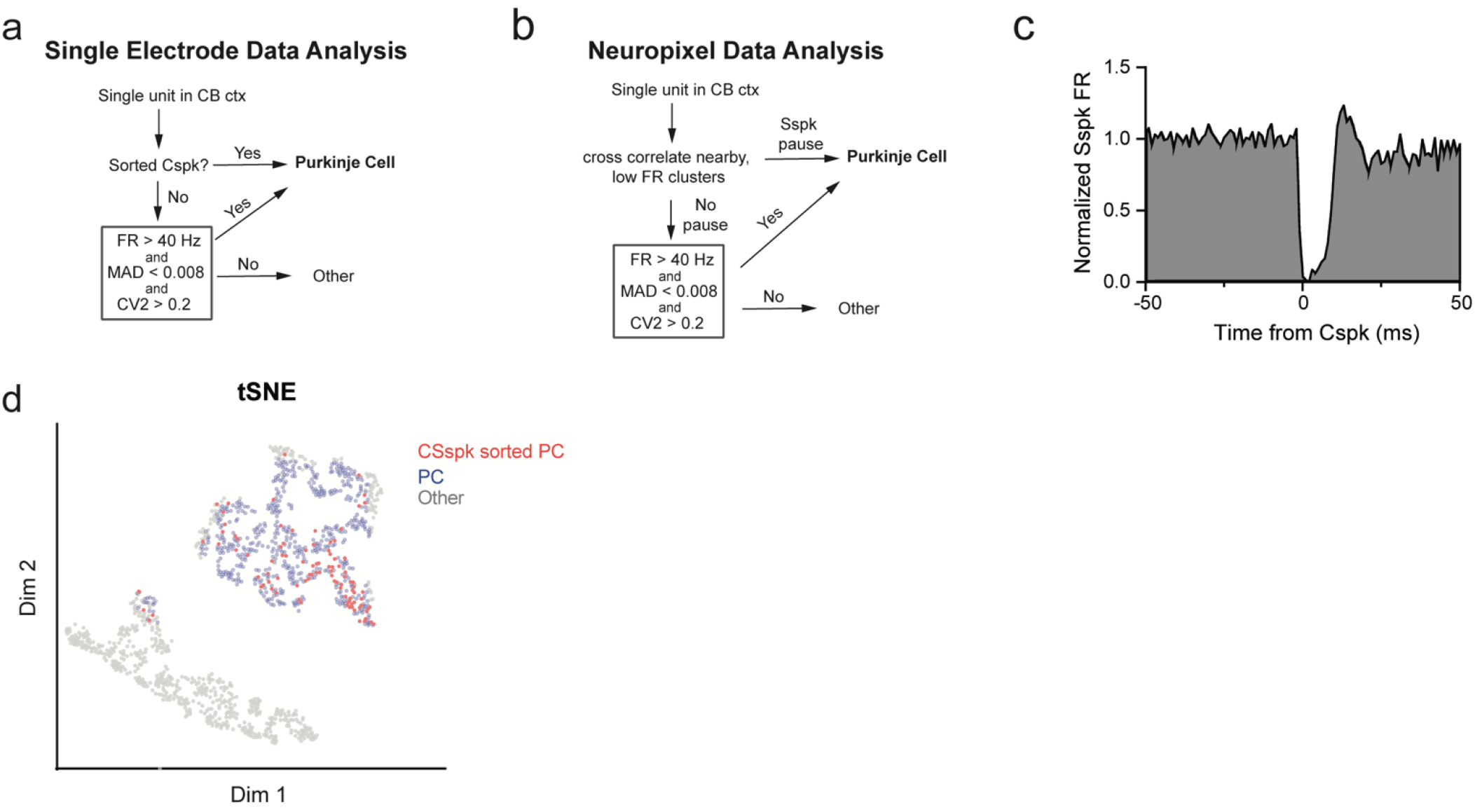
PC identification by firing rate characteristics. a. Cerebellar recordings using single electrodes were first anatomically targeted to cerebellar cortex. If a recorded cell had visible Cspks they were classified as PCs. Otherwise, if cells had a firing rate > 40 Hz, a median absolute difference firing rate from the median interspike interval (MAD) < 0.008, and a CV2 > 0.2, they were classified as PCs^80^. b. Neuropixel-recorded single units were cross correlated with nearby (<200 microns) low firing rate (<5 Hz) single units. If this cross correlation exhibited the characteristic firing rate pause seen in PC simple spikes after a Cspk, these units were classified as the simple spikes and Cspks of a single PC. If no pause was seen, cells that exhibited the same firing rate, MAD, and CV2 profile described in **a** were classified as PCs. c. Example simple spike pause aligned to the time of a Cspk from a Neuropixel recording. d. Embedding MAD, CV2, and FR into a two-dimensional space using tSNE shows two distinct clusters, one corresponding largely to cells that were identified using the criteria in **a** and **b** and the other corresponding to other cells (n = 1268 sorted cells).

**Figure S3.**
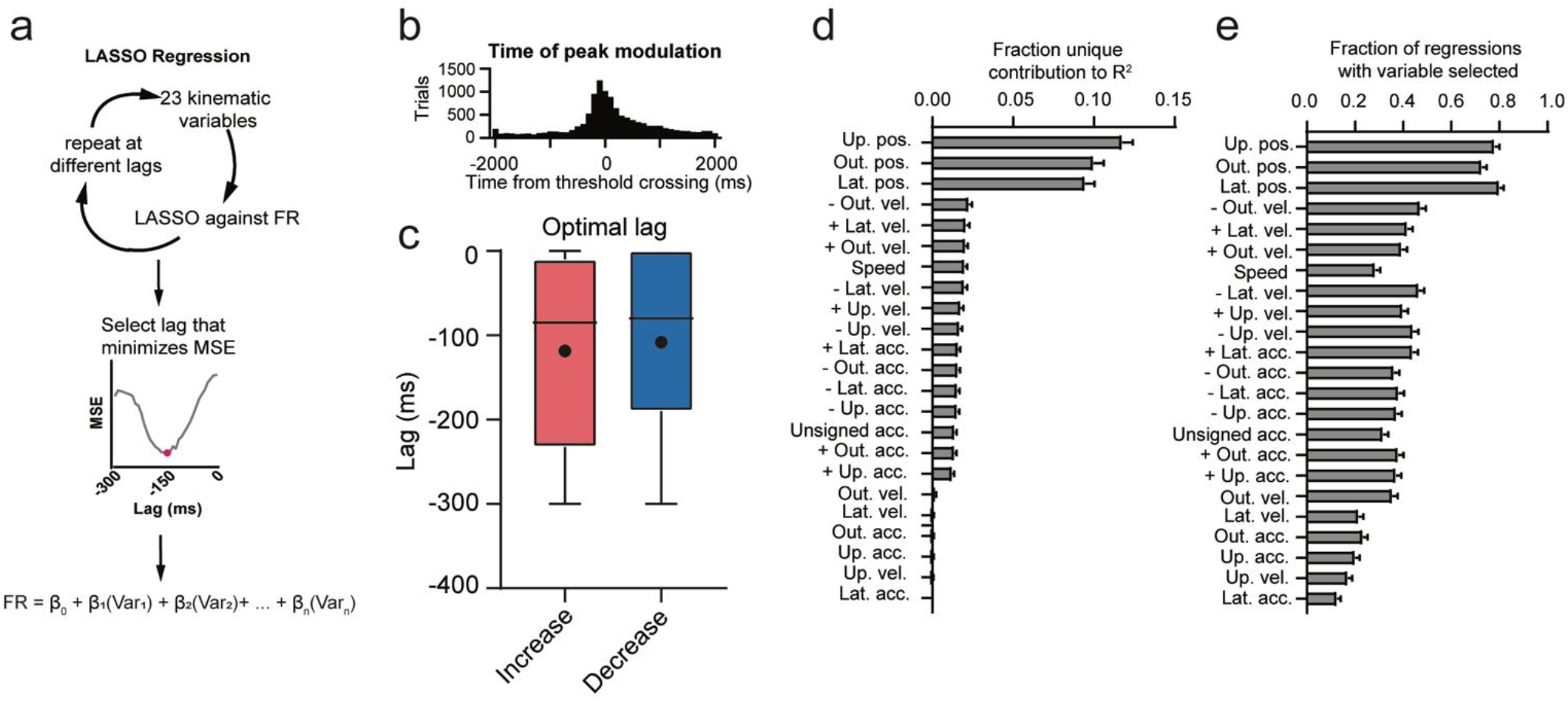
Lasso regression details. a. Schematic of lasso regressions. 23 kinematic variables were regressed against firing rate at different lags from 0 to −300 ms. The lag that minimized the mean squared error (MSE) of the regressions was selected. b. Peak modulation time of all cells across all reaches (n = 11806 trials, 465 cells). c. Optimal lags of the LASSO regression for each cell. d. Fraction of the unique contribution to total variance explained for each regressor. e. Fraction of regressions with each variable selected.

**Figure S4.**
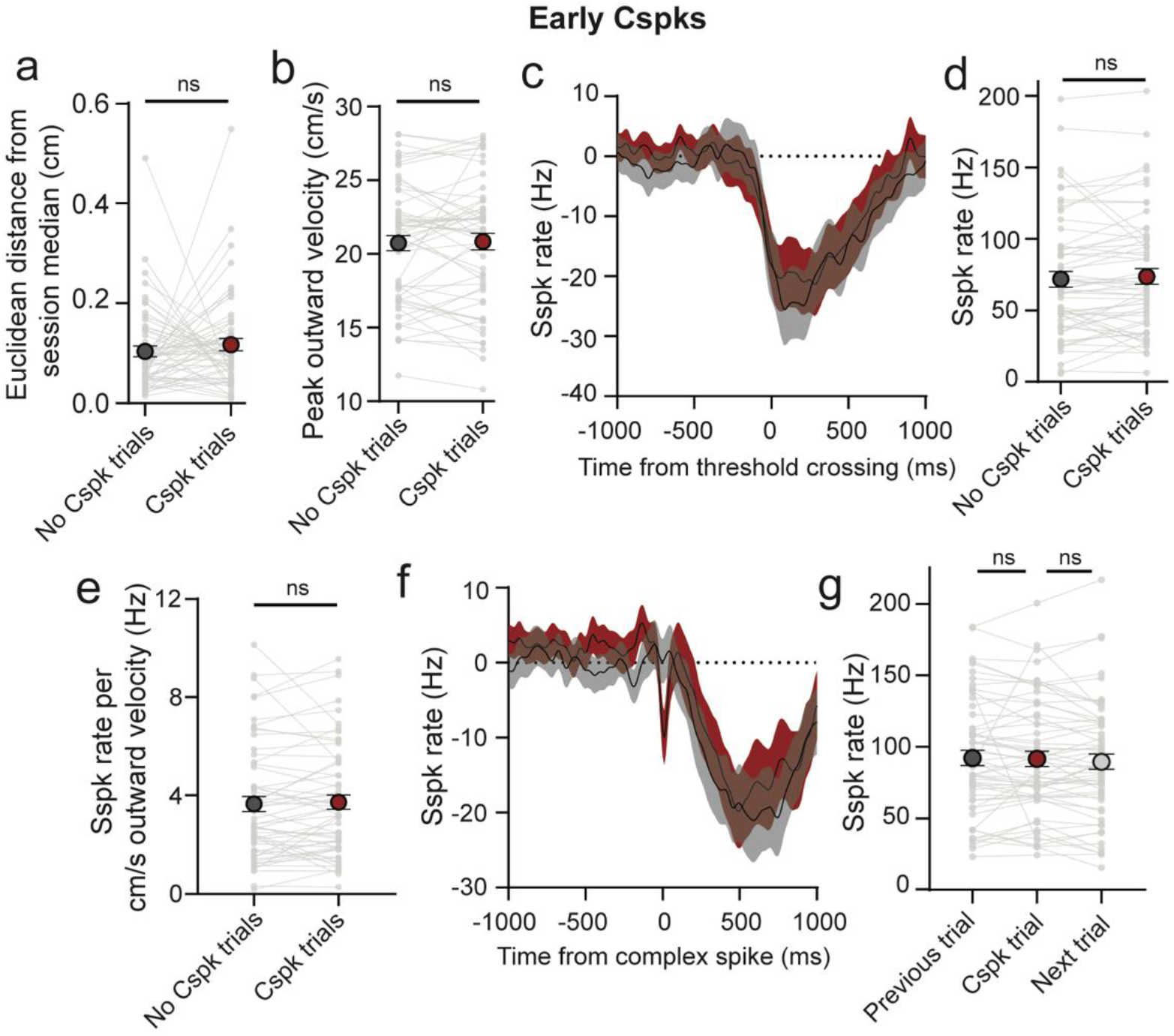
Kinematic and simple spike correlates of early Cspks. a. Cspks in the 500 ms before reach onset were not associated with differences in target error as assessed with euclidean distance form session median compared to non-Cspk trials. b. No difference in peak outward velocity was observed between Cspks and non-Cspk trials. c. Simple spike firing rate in trials with early Cspk and non-Cspk trials. d. No difference in simple spike rate during outreach was seen in early Cspk trials compared with non-Cspk trials. e. No difference in simple spike rate per outward velocity was seen in early Cspk trials compared with non-Cspk trials. f. Simple spike firing aligned to the time of early Cspks compared to similarly aligned trials without early Cspk trials. g. No difference in simple spike rate in the 100 ms preceding early Cspks was seen compared to the similarly aligned previous or next trial.

**Figure S5.**
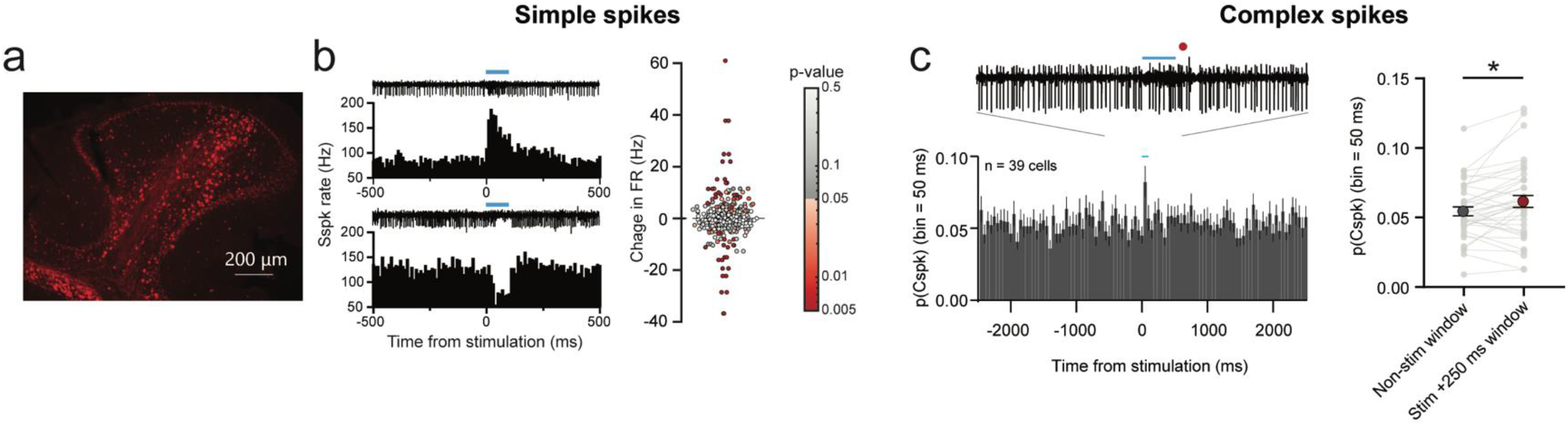
Changes in PC firing during optogenetic stimulation of mossy fibers. a. Mossy fiber boutons expressing hSyn-ChR2-mCherry in the cerebellar cortex. b. Simple spike responses to mossy fiber stimulation. Left: examples of single-cell simple spike responses to mossy fiber stimulation. Right: quantification of simple spike responses to all recorded cells. Significance of differences are indicated by the color and corresponding p-value map. c. Cspk responses to mossy fiber stimulation. Left: PSTH of the population of recorded cells with Cspks binned at 50 ms. A single trace showing a Cspk after stimulation is shown above. Right: Quantification of Cspk probability in the 250 ms after stimulation and non-stimulated epochs for each cell.

**Figure S6.**
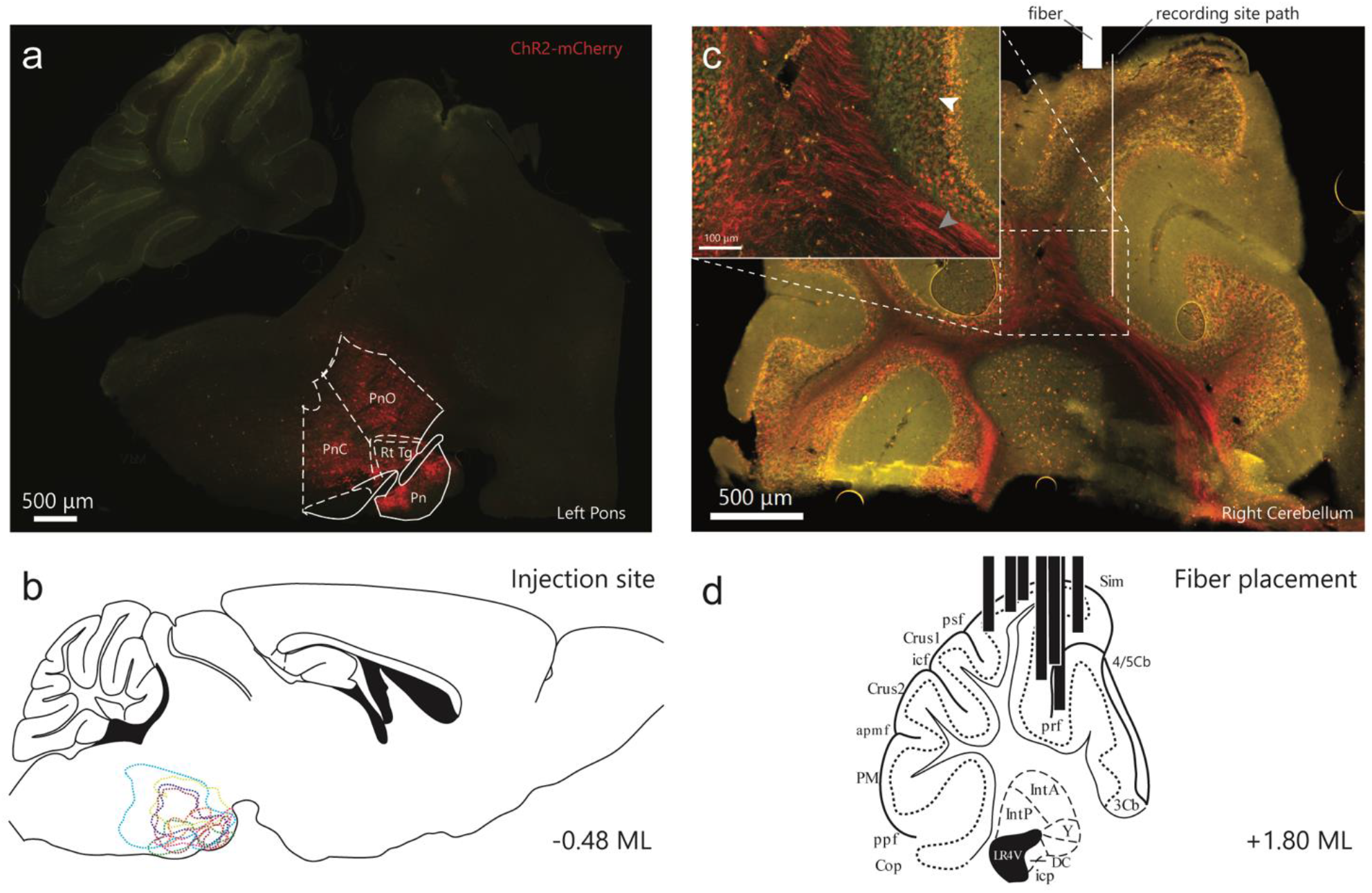
Opsin expression for mice in behavioral experiments. a. Histological section showing ChR2-mCherry expression at the injection site in the left pontine nuclei (Pn: pontine nuclei; RtTg: reticulotegmental nuclei; PnO: pontine reticular nuclei, oral part; PnC: pontine reticular nuclei, caudal part). b. Contours of ChR2 expression in the pontine nuclei for mice used in behavioral experiments. c. Right cerebellum of the animal shown in **a**. Mossy fiber axons (grey arrow) and boutons (white arrow) can be seen expressing ChR2 in the cerebellar cortex. The approximate location of the optical fiber and recording site path are shown in white. d. Location of fiber placement in a representative section for animals used in behavioral experiments.

**Figure S7.**
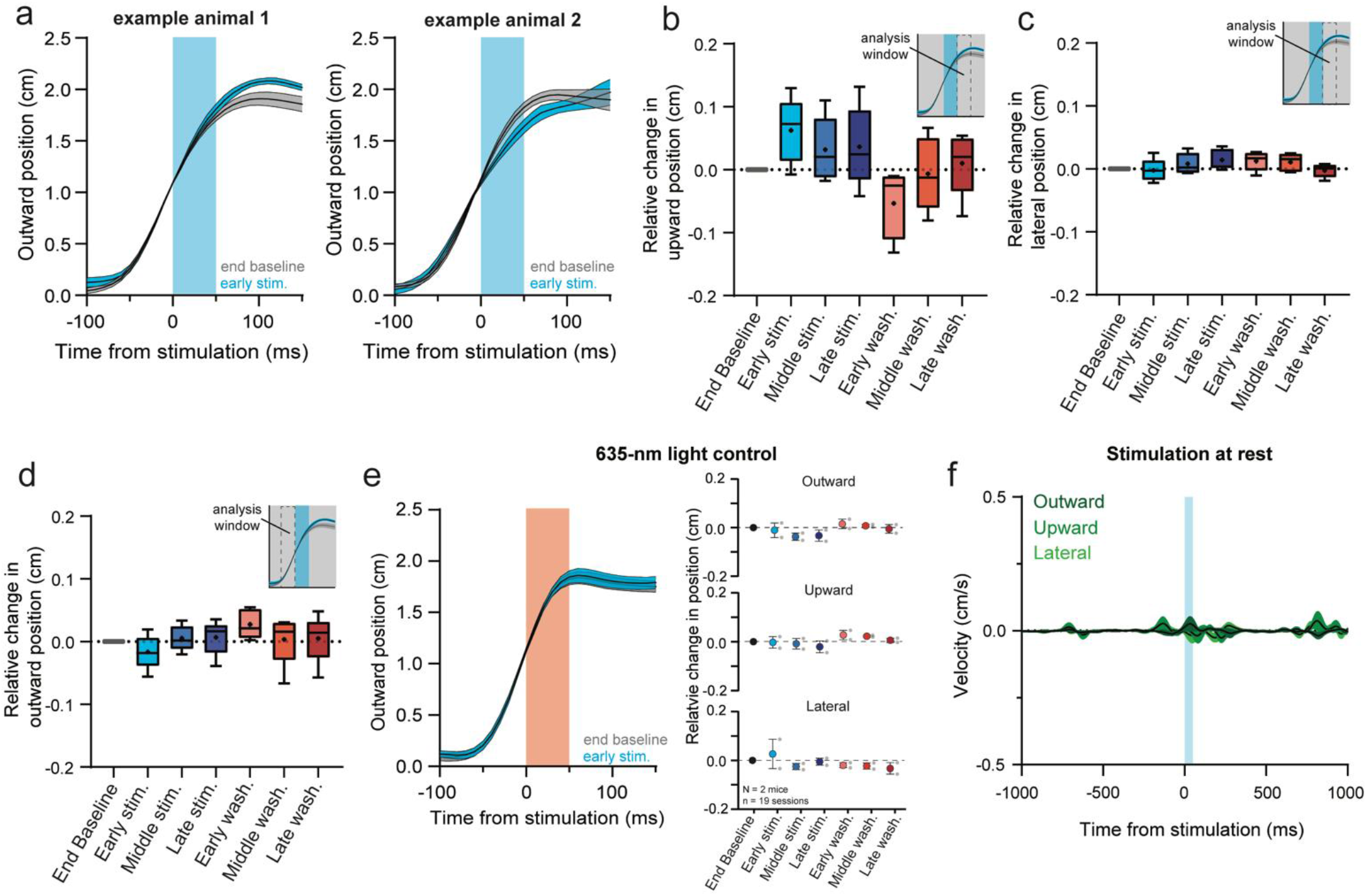
Fixed-position stimulation supplemental data a. Two example mice with differing effects of stimulation on early reaches in the stimulation block. To account for diverging effects we define the direction of deviation with stimulation as positive and the opposing direction as negative. b. Summary of the relative change in upward position for the same data shown in Fig. 3e. Relative change in upward position was assessed in the 50-ms window following the end of stimulation. c. Summary of the relative change in lateral position for the same data shown in Fig. 3e. d. Summary of the relative change in outward position for in the 50-ms window before stimulation. e. Stimulating with 635-nm light did not cause deviations in position or adaptation profiles. f. Stimulating while the mouse was awake with its hand at rest on the bar produced virtually no movement.

**Figure S8.**
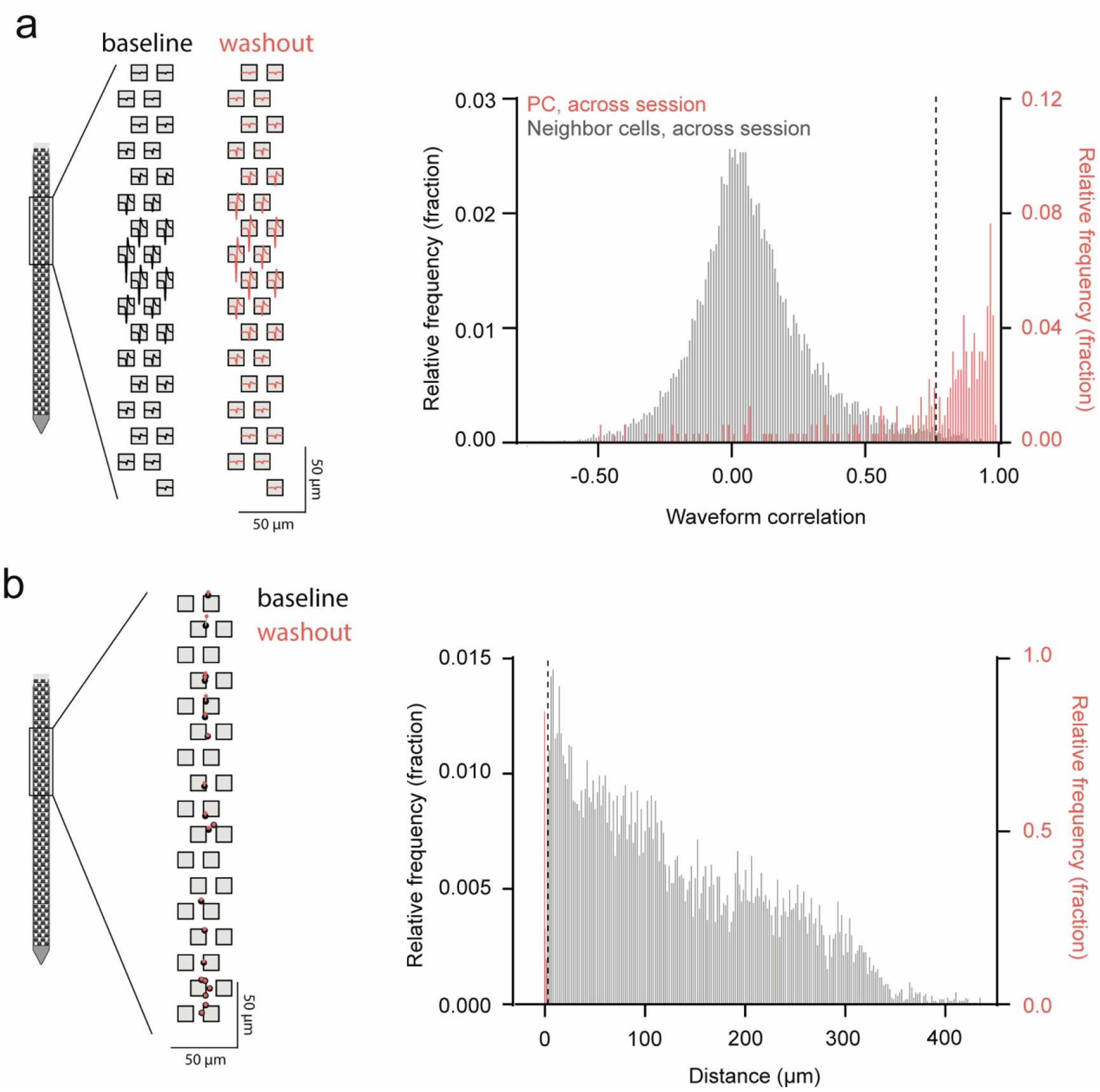
Assessing unit stability across recording sessions. a. Left: Waveforms templates detected on each Neuropixel electrode for a cell during baseline and during washout. Right: Histogram of waveform correlation of PCs across sessions (red) and of mismatched neighboring cells, across the session (shuffled control, grey). PCs with an across-session waveform correlation that fell below the 99^th^ percentile of the shuffled control (dashed line) were excluded from further analysis. b. Left: Unit displacement for cells across a session. Baseline unit position is shown in grey and washout position is shown in red. Right: Histogram of unit displacement of PCs across sessions (red) and of mismatched neighboring cells, across the session (shuffled control, grey). PCs with an across-session displacement that fell below the 1^st^ percentile of the shuffled control (dashed line) were excluded from further analysis.

**Figure S9.**
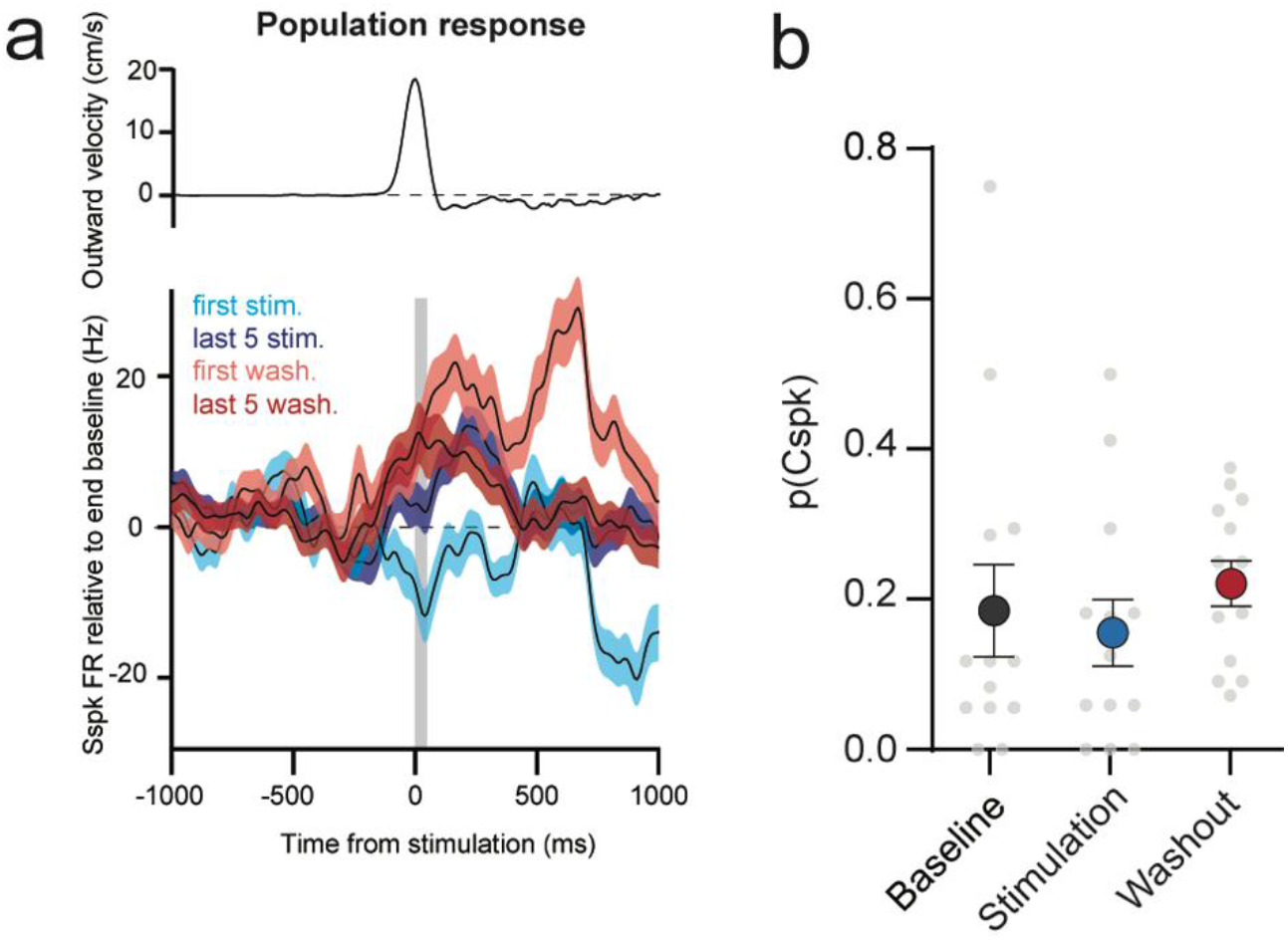
Population simple spike and complex spike responses during fixed-position stimulation experiments. **a.** Same as data shown in Fig. 4g with the last 5 stimulated and washout reaches included. The initial stimulation and washout effects are reduced across the stimulation and washout blocks, respectively. **b.** Cspks analyzed during fixed position stimulation experiments for the baseline, stimulation, and washout blocks (n = 13 cells with sorted Cspks).

**Figure S10.**
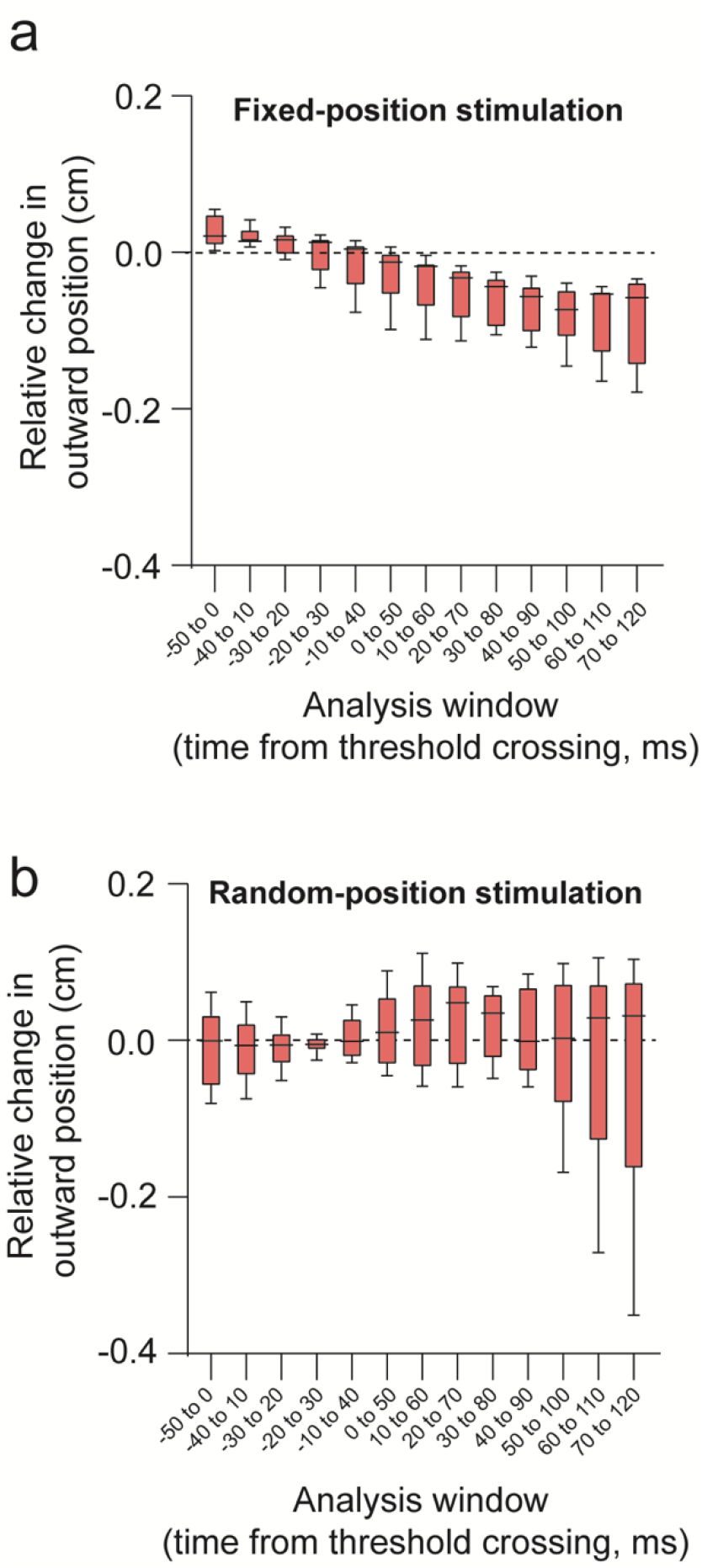
Temporal analysis of early washout effect for fixed-position and random-stimulation experiments. **a.** Analysis of fixed-position stimulation experiment early washout reaches in 50-ms time windows across the reach. Each window is shifted 10 ms from the adjacent time window. Aftereffect emerges around the time stimulation was delivered in the stimulation block. **b.** Same as **a** but for random-position stimulation experiments. Consistent aftereffects relative to baseline reaches do not emerge in any of the analyzed windows.

**Figure S11.**
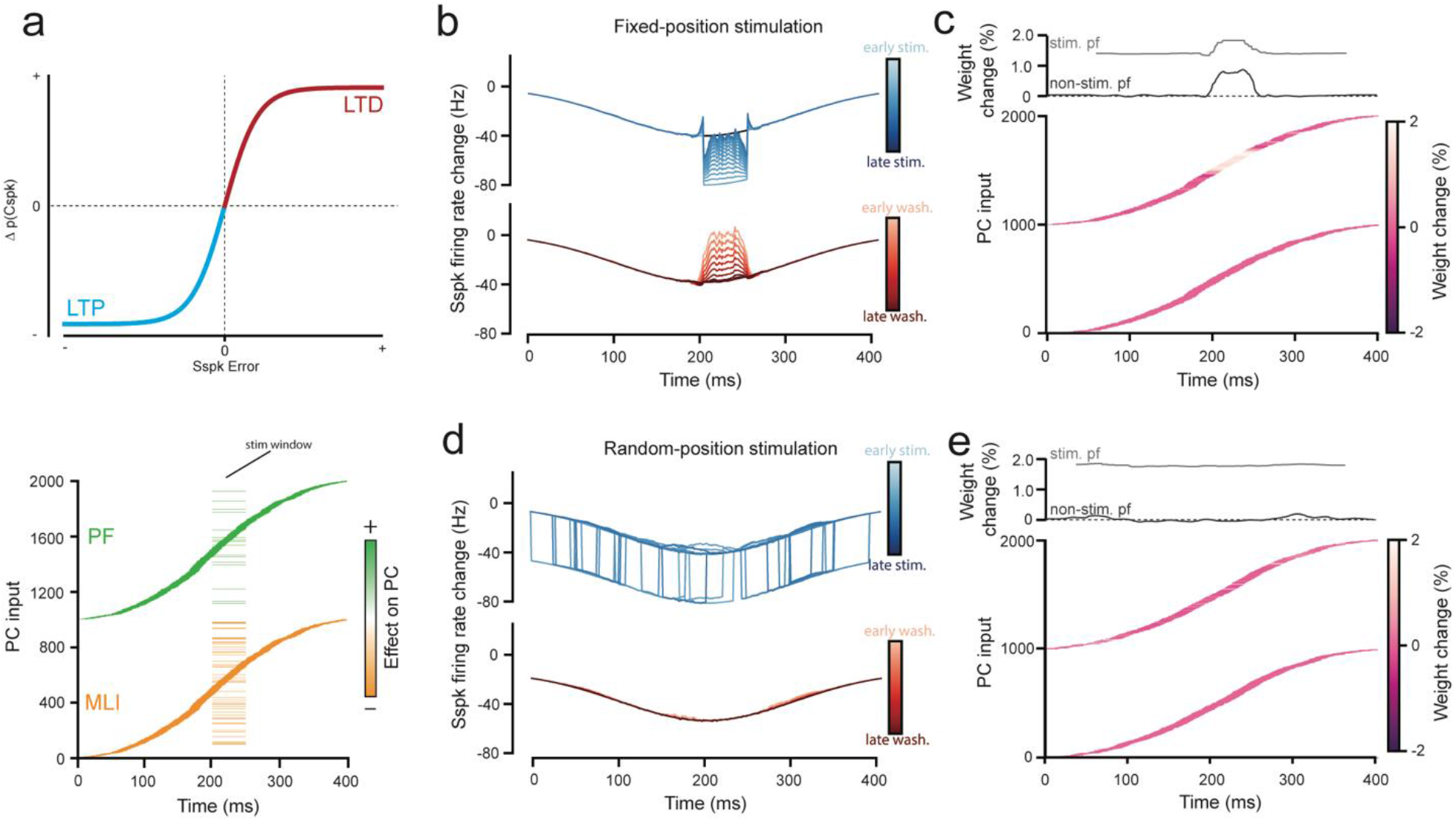
Cerebellar model adaptation to a negative-going perturbation. **a.** Model as described in Fig. 6. In this case the number of stimulated MLIs is greater than the number of parallel fibers (bottom) leading to a net negative stimulation effect. Negative simple spike error lowers the probability of Cspks below baseline, leading to LTP (top). **b.** PC simple spike activity during the stimulation block and washout block of fixed-position stimulation as described in Fig. 6b. Here the stimulation reduces firing rate. **c.** Same as described in Fig. 6c. Here parallel fiber weight changes increase to compensate for the stimulation. Note that while not displayed the quantification of the adaptation is identical to the data displayed in Fig. 6d. **d.** Comparison of model output to empirical observations for fixed-position stimulus conditions (Fig.3). Model closely matches behavioral adaptation. **e.** Same as **b.** but here the stimulation window is randomized across the reach. Note that while not displayed the quantification of the adaptation is identical to the data displayed in Fig. 6d.

## References

1. Holmes, G. The symptoms of acute cerebellar injuries due to gunshot injuries. Brain 40, 461–535(1917).

2. Becker, M. I. & Person, A. L. Cerebellar Control of Reach Kinematics for Endpoint Precision. Neuron 103, 335–348.e5 (2019).

3. Lang, C. E. & Bastian, A. J. Cerebellar subjects show impaired adaptation of anticipatory EMG during catching. J. Neurophysiol. 82, 2108–2119(1999).

4. Baizer, J. S., Kralj-Hans, I. & Glickstein, M. Cerebellar lesions and prism adaptation in macaque monkeys. J. Neurophysiol. 81, 1960–1965(1999).

5. Tseng, Y. W., Diedrichsen, J., Krakauer, J. W., Shadmehr, R. & Bastian, A. J. Sensory prediction errors drive cerebellum-dependent adaptation of reaching. J. Neurophysiol. 98, 54–62(2007).

6. Morton, S. M. & Bastian, A. J. Cerebellar contributions to locomotor adaptations during splitbelt treadmill walking. J. Neurosci. 26, 9107–9116(2006).

7. Smith, M. A. & Shadmehr, R. Intact ability to learn internal models of arm dynamics in Huntington’s disease but not cerebellar degeneration. J. Neurophysiol. 93, 2809–2821(2005).

8. Miall, R. & Wolpert, D. Forward Models for phyiological motor control. Neural Networks 9, 1265–1279(1996).

9. Kawato, M. & Gomi, H. The cerebellum and VOR / OKR learning models. Trends Neurosci. 15, 445–453(1992).

10. Ohyama, T., Nores, W. L., Murphy, M. & Mauk, M. D. What the cerebellum computes. Trends Neurosci. 26, 222–227(2003).

11. Shadmehr, R. & Krakauer, J. W. A computational neuroanatomy for motor control. Exp. Brain Res. 185, 359–381(2008).

12. Shadmehr, R., Smith, M. A. & Krakauer, J. W. Error correction, sensory prediction, and adaptation in motor control. Annu. Rev. Neurosci. 33, 89–108(2010).

13. Lee, K. H. et al. Circuit mechanisms underlying motor memory formation in the cerebellum. Neuron 86, 529–540(2015).

14. Ekerot, C. F., Garwicz, M. & Schouenborg, J. Topography and nociceptive receptive fields of climbing fibres projecting to the cerebellar anterior lobe in the cat. J. Physiol. 441, 257–274(1991).

15. Bloedel, J. R. & Bracha, V. On the cerebellum, cutaneomuscular reflexes, movement control and the elusive engrams of memory. Behav. Brain Res. 68, 1–44(1995).

16. Lavond, D. G., Knowlton, B. J., Steinmetz, J. E. & Thompson, R. F. Classical Conditioning of the Rabbit Eyelid Response With a Mossy-Fiber Stimulation CS: II. Lateral Reticular Nucleus Stimulation. Behav. Neurosci. 101, 676–682(1987).

17. Mauk, M. D., Steinmetz, J. E. & Thompson, R. F. Classical conditioning using stimulation of the inferior olive as the unconditioned stimulus. Proc. Natl. Acad. Sci. U. S. A. 83, 5349–5353(1986).

18. Coesmans, M., Weber, J. T., De Zeeuw, C. I. & Hansel, C. Bidirectional parallel fiber plasticity in the cerebellum under climbing fiber control. Neuron 44, 691–700(2004).

19. Ito, M. & Kano, M. Long-lasting depression of parallel fiber-Purkinje cell transmission induced by conjunctive stimulation of parallel fibers and climbing fibers in the cerebellar cortex. Neurosci. Lett. 33, 253–258(1982).

20. Ito, M. Cerebellar control of the vestibulo-ocular reflex--around the flocculus hypothesis. Annu. Rev. Neurosci. 5, 275–297(1982).

21. Blazquez, P. M., Hirata, Y., Heiney, S. A., Green, A. M. & Highstein, S. M. Cerebellar Signatures of Vestibulo-Ocular Reflex Motor Learning. J. Neurosci. 23, 9742–9751(2003).

22. Rowan, M. J. M. et al. Graded Control of Climbing-Fiber-Mediated Plasticity and Learning by Inhibition in the Cerebellum. Neuron 1–17 (2018). doi:10.1016/j.neuron.2018.07.024

23. Yang, Y. & Lisberger, S. G. Interaction of plasticity and circuit organization during the acquisition of cerebellum-dependent motor learning. Elife 2013, 1–19(2013).

24. Herzfeld, D. J., Kojima, Y., Soetedjo, R. & Shadmehr, R. Encoding of error and learning to correct that error by the Purkinje cells of the cerebellum. Nat. Neurosci. 21, (2018).

25. Raymond, J. L., Lisberger, S. G. & Mauk, M. D. The cerebellum: A neuronal learning machine? Science (80-.). 272, 1126–1131(1996).

26. Mathis, M. W., Mathis, A. & Uchida, N. Somatosensory Cortex Plays an Essential Role in Forelimb Motor Adaptation in Mice. Neuron 93, 1493–1503.e6 (2017).

27. Ito, M. Neural design of the cerebellar motor control system. Brain Res. 40, 81–84(1972).

28. Weiler, J., Gribble, P. L. & Pruszynski, J. A. Spinal stretch reflexes support efficient hand control. Nat. Neurosci. 22, 529–533(2019).

29. Gandolfo, F., Li, C. S. R., Benda, B. J., Padoa Schioppa, C. & Bizzi, E. Cortical correlates of learning in monkeys adapting to a new dynamical environment. Proc. Natl. Acad. Sci. U. S. A. 97, 2259–2263(2000).

30. Li, C. S. R., Padoa-Schioppa, C. & Bizzi, E. Neuronal correlates of motor performance and motor learning in the primary motor cortex of monkeys adapting to an external force field. Neuron 30, 593–607(2001).

31. Proville, R. D. et al. Cerebellum involvement in cortical sensorimotor circuits for the control of voluntary movements. Nat. Neurosci. 17, 1233–1239(2014).

32. Hewitt, A. L., Popa, L. S. & Ebner, T. J. Changes in Purkinje Cell Simple Spike Encoding of Reach Kinematics during Adaption to a Mechanical Perturbation. J. Neurosci. 35, 1106–1124(2015).

33. Becker, M. I., Calame, D. J., Wrobel, J. & Person, A. L. Online control of reach accuracy in mice. J. Neurophysiol. 124, 1637–1655(2020).

34. Low, A. Y. T. et al. Precision of Discrete and Rhythmic Forelimb Movements Requires a Distinct Neuronal Subpopulation in the Interposed Anterior Nucleus. Cell Rep. 22, 2322–2333(2018).

35. Guo, J.-Z. et al. Disrupting cortico-cerebellar communication impairs dexterity. Elife 10, 1–31(2021).

36. Tibshirani, R. Regression Shrinkage and Selection via the Lasso. J. R. Stat. Soc. 58, 267–288(1996).

37. Popa, L. S., Streng, M. L. & Ebner, T. J. Long-Term Predictive and Feedback Encoding of Motor Signals in the Simple Spike Discharge of Purkinje Cells. 4, (2017).

38. Streng, M. L., Popa, L. S. & Ebner, T. J. Climbing Fibers Control Purkinje Cell Representations of Behavior. J. Neurosci. 37, 1997–2009(2017).

39. Coltz, J. D., Johnson, M. T. & Ebner, T. J. Cerebellar Purkinje cell simple spike discharge encodes movement velocity in primates during visuomotor arm tracking. J. Neurosci. 19, 1782–803(1999).

40. Hewitt, A. L., Popa, L. S., Pasalar, S., Hendrix, C. M. & Ebner, T. J. Representation of limb kinematics in Purkinje cell simple spike discharge is conserved across multiple tasks. J. Neurophysiol. 106, 2232–2247(2011).

41. Roitman, A. V., Pasalar, S., Johnson, M. T. V. & Ebner, T. J. Position, direction of movement, and speed tuning of cerebellar Purkinje cells during circular manual tracking in monkey. J. Neurosci. 25, 9244–9257(2005).

42. Musall, S., Kaufman, M. T., Juavinett, A. L., Gluf, S. & Churchland, A. K. Single-trial neural dynamics are dominated by richly varied movements. Nat. Neurosci. 22, 1677–1686(2019).

43. Hewitt, A. L., Popa, L. S. & Ebner, T. J. Changes in Purkinje Cell Simple Spike Encoding of Reach Kinematics during Adaption to a Mechanical Perturbation. J. Neurosci. 35, 1106–1124(2015).

44. Ebner, T. J., Hewitt, A. L. & Popa, L. S. What Features of Limb Movements are Encoded in the Discharge of Cerebellar Neurons? The Cerebellum 10, 683–693(2010).

45. Herzfeld, D. J., Kojima, Y., Soetedjo, R. & Shadmehr, R. Encoding of action by the Purkinje cells of the cerebellum. Nature 526, 439–441(2015).

46. Brown, S. T. & Raman, I. M. Sensorimotor Integration and Amplification of Reflexive Whisking by Well-Timed Spiking in the Cerebellar Corticonuclear Circuit. Neuron 1–12 (2018). doi:10.1016/j.neuron.2018.06.028

47. Person, A. L. & Raman, I. M. Purkinje neuron synchrony elicits time-locked spiking in the cerebellar nuclei. Nature 481, 502–505(2012).

48. Thier, P., Dicke, P. W., Haas, R. & Barash, S. Encoding of movement time by populations of cerebellar Purkinje cells. Nature 405, 72–76(2000).

49. Medina, J. F. & Lisberger, S. G. Links from complex spikes to local plasticity and motor learning in the cerebellum of awake-behaving monkeys. Nat. Neurosci. 11, 1185–1192(2008).

50. Kimpo, R. R., Rinaldi, J. M., Kim, C. K., Payne, H. L. & Raymond, J. L. Gating of neural error signals during motor learning. Elife 3, 1–23(2014).

51. Avraham, G., Taylor, J. A., Breska, A., Ivry, R. B. & McDougle, S. D. Contextual effects in motor adaptation adhere to associative learning rules. bioRxiv 2020.09.14.297143 (2021).

52. Wagner, M. J. et al. A neural circuit state change underlying skilled movements. Cell 3731–3747 (2021). doi:10.1016/j.cell.2021.06.001

53. Kitazawa, S., Kimura, T. & Yin, P. B. Cerebellar complex spikes encode both destinations and errors in arm movements. Nature 392, 494–497(1998).

54. Gaffield, M. A. & Christie, J. M. The cerebellum encodes and influences the initiation and termination of discontinuous movements. bioRxiv 1–31 (2021).

55. Chabrol, F. P., Blot, A. & Mrsic-Flogel, T. D. Cerebellar Contribution to Preparatory Activity in Motor Neocortex. Neuron 103, 506–519.e4 (2019).

56. Ekerot, C.-F. & Kano, M. Long-term depression of parallel fibre synapses following stimulation of climbing fibres. Brain Res. 342, 357–360(1985).

57. Ito, M., Sakurai, M. & Tongroach, P. Climbing fibre induced depression of both mossy fibre responsiveness and glutamate sensitivity of cerebellar Purkinje cells. J. Physiol. 324, 113–134(1982).

58. Miall, R. C., Keating, J. G., Malkmus, M. & Thach, W. T. Simple spike activity predicts occurrence of complex spikes in cerebellar Purkinje cells. Nat. Neurosci. 1, 13–15(1998).

59. Brodal, A. & Jansen, J. The ponto-cerebellar projection in the rabbit and cat. J. Comp. Neurol. 84, 31–118(1946).

60. Brodal, P. & Bjaalie, J. G. Organization of the pontine nuclei. Neurosci. Res. 13, 83–118(1992).

61. Leergaard, T. B. et al. Three-dimensional topography of corticopontine projections from rat sensorimotor cortex: Comparisons with corticostriatal projections reveal diverse integrative organization. J. Comp. Neurol. 478, 306–322(2004).

62. Sillitoe, R. V., Fu, Y. H. & Watson, C. Cerebellum. The Mouse Nervous System (Elsevier Inc., 2012). doi:10.1016/B978-0-12-369497-3.10011-1

63. Huang, C. C. et al. Convergence of pontine and proprioceptive streams onto multimodal cerebellar granule cells. Elife 2013, 1–17(2013).

64. Kim, J. & Augustine, G. J. Molecular Layer Interneurons: Key Elements of Cerebellar Network Computation and Behavior. Neuroscience 462, 22–35(2021).

65. Rasmussen, A., Jirenhed, D. A., Wetmore, D. Z. & Hesslow, G. Changes in complex spike activity during classical conditioning. Front. Neural Circuits 8, 1–13(2014).

66. Herzfeld, D. J., Hall, N. J., Tringides, M. & Lisberger, S. G. Principles of operation of a cerebellar learning circuit. Elife 9, 1–30(2020).

67. Mauk, M. D. & Buonomano, D. V. The neural basis of temporal processing. Annu. Rev. Neurosci. 27, 307–340(2004).

68. Shimansky, Y., Wang, J. J., Bauer, R. A., Bracha, V. & Bloedel, J. R. On-line compensation for perturbations of a reaching movement is cerebellar dependent: Support for the task dependency hypothesis. Exp. Brain Res. 155, 156–172(2004).

69. Albergaria, C., Silva, N. T., Pritchett, D. L. & Carey, M. R. Locomotor activity modulates associative learning in mouse cerebellum. Nat. Neurosci. 21, 725–735(2018).

70. Donchin, O., Francis, J. T. & Shadmehr, R. Quantifying generalization from trial-by-trial behavior of adaptive systems that learn with basis functions: Theory and experiments in human motor control. J. Neurosci. 23, 9032–9045(2003).

71. Nashef, A. et al. Cerebellar Shaping of Motor Cortical Firing Is Correlated with Timing of Motor Actions Article Cerebellar Shaping of Motor Cortical Firing Is Correlated with Timing of Motor Actions. CellReports 23, 1275–1285(2018).

72. Shinoda, Y., Kakei, S., Futami, T. & Wannier, T. Thalamocortical organization in the cerebello-thalamo-cortical system. Cereb. Cortex 3, 421–429(1993).

73. Shinoda, Y., Futami, T. & Kano, M. Input-output organization of the ventrolateral nucleus of the thalamus. Stereotact Funct Neurosurg 60, 17–31(1993).

74. Shinoda, Y., Futami, T. & Kano, M. Synaptic organization of the cerebello-thalamo-cerebral pathway in the cat. II. Input-output organization of single thalamocortical neurons in the ventrolateral thalamus. Neurosci. Res. 2, 157–180(1985).

75. White, J. J. et al. An optimized surgical approach for obtaining stable extracellular single-unit recordings from the cerebellum of head-fixed behaving mice. J. Neurosci. Methods 262, 21–31(2016).

76. TestArduino. (2015).

77. Yu, B., Gabriel, D., Noble, L. & An, K. N. Estimate of the optimum cut-off frequency for the Butterworth low-pass digital filter. J. Appl. Biomech. 15, 318–329(1999).

78. Sedaghat-Nejad, E. et al. P-sort: An open-source software for cerebellar neurophysiology. J. Neurophysiol. 126, 1055–1075(2021).

79. Pachitariu, M., Steinmetz, N., Kadir, S., Carandini, M. & Kenneth D. H., Kilosort: realtime spike-sorting for extracellular electrophysiology with hundreds of channels. bioRxiv 061481 (2016). doi:10.1101/061481

80. Hensbroek, R. A. et al. Identifying Purkinje cells using only their spontaneous simple spike activity. J. Neurosci. Methods 232, 173–180(2014).

81. Schoonover, C. E., Ohashi, S. N., Axel, R. & Fink, A. J. P. Representational drift in primary olfactory cortex. Nature 594, 541–546(2021).

82. Kennedy, A. et al. A temporal basis for predicting the sensory consequences of motor commands in an electric fish. Nat. Neurosci. 17, 416–422(2014).

83. Lanore, F., Cayco-Gajic, N. A., Gurnani, H., Coyle, D. & Silver, R. A. Cerebellar granule cell axons support high-dimensional representations. Nat. Neurosci. 24, 1142–1150(2021).

84. Wagner, M. J. et al. Shared cortex-cerebellum dynamics in the execution and learning of a motor task. Cell 177, 669–682(2019).

85. Jörntell, H. & Hansel, C. Synaptic memories upside down: bidirectional plasticity at cerebellar parallel fiber-Purkinje cell synapses. Neuron 52, 227–238(2006).

